# Spatial modelling for population replacement of mosquito vectors at continental scale

**DOI:** 10.1101/2021.10.06.463299

**Authors:** Nicholas J. Beeton, Andrew Wilkins, Adrien Ickowicz, Keith R. Hayes, Geoffrey R. Hosack

## Abstract

Malaria is one of the deadliest vector-borne diseases in the world. Researchers are developing new genetic and conventional vector control strategies to attempt to limit its burden. Novel control strategies require detailed safety assessment to ensure responsible and successful deployments. *Anopheles gambiae* sensu stricto (s.s.) and *Anopheles coluzzii*, two closely related subspecies within the species complex *Anopheles gambiae* sensu lato (s.l.), are among the dominant malaria vectors in sub-Saharan Africa. These two subspecies readily hybridise and compete in the wild and are also known to have distinct niches, each with spatially and temporally varying carrying capacities driven by precipitation and land use factors.

We model the spread and persistence of a population-modifying gene drive system in these subspecies across sub-Saharan Africa by simulating introductions of genetically modified mosquitoes across the African mainland and its offshore islands. We explore transmission of the gene drive between the two subspecies that arise from different hybridisation mechanisms, the effects of both local dispersal and potential wind-aided migration to the spread, and the development of resistance to the gene drive. Given the best current available knowledge on the subspecies’ life histories, we find that an introduced gene drive system with typical characteristics can plausibly spread from even distant offshore islands to the African mainland with the aid of wind-driven migration, with resistance taking over within a decade. Our model accounts for regional to continental scale mechanisms, and demonstrates a range of realistic dynamics including the effect of prevailing wind on spread and spatio-temporally varying carrying capacities for subspecies. As a result, it is well-placed to answer future questions relating to mosquito gene drives as important life history parameters become better understood.

## 1 Introduction

Contemporary malaria control interventions – insecticide treated bed nets, indoor residual spraying and artemisinin based combination therapy – have dramatically reduced the burden of malaria in Africa (Bhatt et al., 2015). Since 2017, however, the rate of progress on malaria reduction has stalled and in 2019 malaria still claimed an estimated 389,000 African lives, mainly children under 5 years of age (World Health Organisation, 2020). At least 99% of these cases are caused by *Plasmodium falciparum*, transmitted by a small number of dominant malaria vectors, most notably *Anopheles arabiensis, Anopheles coluzzii, Anopheles gambiae* sensu stricto (s.s.) and *Anopheles funestus* (Wiebe et al., 2017).

The ongoing burden of malaria, together with increasing rates of insecticide resistance in malarial vector mosquitoes (Wiebe et al., 2017), has motivated proposals to develop new genetic control strategies, including: a) self-limiting, population suppression methods that induce male sterility (Windbichler et al., 2008; Klein et al., 2012) or male bias (Galizi et al., 2014; Facchinelli et al., 2019); b) self-sustaining (gene drive), suppression methods that induce female sterility (Hammond et al., 2016; Kyrou et al., 2018); and, c) self-sustaining, population replacement methods that make vectors refractory for the malaria parasite (Gantz et al., 2015; Carballar-Lejarazú et al., 2020). Any proposal to test these genetic control strategies outside of contained laboratory settings will likely require a detailed quantitative risk assessment that predicts the potential spread and persistence of transgenic mosquitoes from release sites, and the possible introgression of a transgenic construct into closely related species through interspecific mating (James et al., 2018, 2020). Spatial models of spread and persistence are also needed to describe the dynamics of important gene drive processes such as the development of resistance to the gene drive (Price et al., 2020). Quantitative spatial models have been developed for the spread and persistence of self-limiting, population suppressing constructs (Dufourd and Dumont, 2013; Facchinelli et al., 2019), together with self-sustaining, population-modifying (Beaghton et al., 2017; Tanaka et al., 2017; Noble et al., 2018; Wyse et al., 2018) and population-suppressing (Eckhoff et al., 2016; North and Godfray, 2018; North et al., 2019) constructs.

This paper models the spread and persistence of a population-modifying gene drive system (Alphey et al., 2020; Nolan, 2021) in *Anopheles gambiae* s.s. and *Anopheles coluzzii* across sub-Saharan Africa. These two subspecies, which are often modelled as single group, are together with *Anopheles arabiensis* and *Anopheles funestus* the dominant vectors of malaria in sub-Saharan Africa. *An. gambiae* s.s. and *An. coluzzii* are both highly anthropophilic and efficient malaria vectors. The two subspecies are closely related enough to interbreed but hybridisation rates vary in space and time (Pombi et al., 2017). They also have different larval habitat preferences (Lehmann and Diabate, 2008), and *An. coluzzii* is thought to have a superior resistance to desiccation stress (Tene Fossog et al., 2015), hence is more drought tolerant than *An. gambiae* s.s. (Sinka et al., 2016). Although they are defined as two separate species by Coetzee et al. (2013), we refer to them here and subsequently as subspecies to emphasise the lack of reproductive isolation between the two taxa (see Pombi et al., 2017; Tene Fossog et al., 2015), which is a focus of our model.

In addition to general concerns for gene drives such as the development of resistance, the following ecological hypotheses proposed in the literature are investigated: transmission of the gene drive between two hybridising subspecies of *Anopheles gambiae* sensu lato (s.l.) by vertical gene transfer (Pombi et al., 2017; Beeton et al., 2020; Selvaraj et al., 2020); possible long range dispersal or long distance migration (North and Godfray, 2018; Huestis et al., 2019); and the nature of spatially and temporally varying carrying capacities driven by precipitation and land use factors (White et al., 2011; Tene Fossog et al., 2015; North and Godfray, 2018). This model is designed to provide scenario based testing of structural hypotheses that formalise the current state of knowledge for key gene drive and population life history parameters.

Each of these structural issues are described in the following subsections:

### 1.1 Choice of Construct

Our focus is on the spread and persistence of a gene drive system with near-neutral fitness that incorporates an unavoidable small reproductive payload cost of expression of the nuclease (see Hammond et al. (2016) and Beaghton et al. (2017)) through a spatially and temporally dynamic population with differential gene flow across sub-Saharan Africa. This scenario thereby evaluates the behaviour of an idealised nearly fitness-neutral population replacement gene drive system at the continental scale. Our analysis focusses on three alleles: the wild– type, the genetic construct for a population replacement gene drive and a resistant allele. Together these form a minimal gene drive spatial model (see Beaghton et al., 2017). Further, we assume that the gene drive is activated in a single locus in each parent’s genetic code as in Beaghton et al. (2017). However, we avoid modelling a gene drive “payload” of a nuclease or effector gene as done in their paper. That is, the nuclease and effector gene may be considered as the same unit, with the effector gene either absent or nearly fitness neutral. In practice, an effector would also be likely to exhibit a genetic load on the receiving organism (e.g. Beaghton et al., 2017). Therefore the model predictions are deliberately optimistic in terms of the magnitude of spread and persistence of the construct, and provide an indication of the spread of a population replacement drive for an idealised effector with negligible fitness cost.

### 1.2 Taxonomic Resolution

We model an intervention where the genetic construct has been introgressed into locally sourced, wildtype *An. gambiae* s.s. or *An. coluzzii* mosquitoes, and subsequently released back into this local population. Depending on geographic location, these subspecies of the *An. gambiae* s.l. complex may introgress with each other and potentially other subspecies such as *An. arabiensis* (*Anopheles gambiae* 1000 Genomes Consortium, 2017; Pombi et al., 2017; Clarkson et al., 2020). Studies at the scale of sub-Saharan Africa often do not discriminate between *An. gambiae* s.s. and *An. coluzzii*. For example, Sinka et al. (2012) combined these two subspecies when plotting species distribution maps due to lack of data; *An. arabiensis*, however, was plotted separately. Similarly, North and Godfray (2018) argued that currently there is a lack of available data for parametrising alternative life history strategies of *An. gambiae* s.s. and *An. coluzzii*, and so did not discriminate between these subspecies in their process model. Indeed, we are currently unaware of any analysis of genetic vector control strategies at this continental scale, with explicit spatial and temporal dynamics, that discriminates between these two co-dominant malaria vectors.

We include alternative subspecies in our model because this leads to altered population dynamics via interspecific mating and density dependence effects (Beeton et al., 2020). Here we explore two different approaches to species assignment of first generation hybrids: 1) species assignment by maternal descent, and 2) equal proportions.

### 1.3 Larval carrying capacity

To parameterise a spatial model that discriminates between *An. gambiae* s.s. and *An. coluzzii*, and their inter– and intra–specific density dependence at the larval stage, we require spatially explicit carrying capacity information about each subspecies, which is anticipated to depend on environmental and social covariates. As observation data is relatively sparse at the subspecies level, we approach the problem in two parts. First, we model the larval carrying capacity of the two subspecies taken together using a functional form. Second, we use empirical relative abundance data to spatially model the relative carrying capacities between subspecies.

#### 1.3.1 Total abundance

The carrying capacity of *Anopheles* species in Africa is often expressed as a function of rainfall. For example, White et al. (2011) found exponentially weighting the past 4 days of rainfall gave the best fit when modelling the abundance of *An. gambiae* s.s. and *An. arabiensis* in Nigeria, an approach subsequently adopted by Wu et al. (2020); whilst Magombedze et al. (2018) used a 7 day moving average of rainfall to model the carrying capacity of the aquatic population of *An. gambiae* s.s., *An. coluzzii* and *An. arabiensis* in Mali.

More complex functional forms invoke additional parameters such as the location (and sometimes length or size) of perennial, intermittent, permanent or human-associated water bodies, as in the models developed by Eckhoff (2011) and Lunde et al. (2013). The most relevant approach for our purposes, however, is that of North and Godfray (2018), who group our two proposed subspecies *An. gambiae* s.s. and *An. coluzzii* in a spatially explicit, individual-based model, across an area of West Africa that exhibits significant environmental variation. They predict local larval carrying capacity based on rainfall, as well as level of access to temporary and permanent water courses. We adapt their results for our model of total carrying capacity for the aggregate of *An. coluzzii* and *An. gambiae* s.s.

#### 1.3.2 Relative abundance

*An. coluzzii* has only relatively recently been described as its own subspecies (Coetzee et al., 2013) after its earlier description as a molecular form within *An. gambiae* s.s. (della Torre et al., 2001, 2002). Despite this taxonomic difficulty, several papers have examined differences in habitat preference between *An. gambiae* s.s. and *An. coluzzii*. In particular, Lehmann and Diabate (2008) note that larval predation and competition has led to selection for temporary freshwater habitats in *An. gambiae* s.s. and conversely permanent habitats for *An. coluzzii*. They suggest that this leads to humidity and/or rainfall clines in relative abundance. Other sources provide data suggesting this is the case for rainfall (Touré et al., 1998; Wondji et al., 2005; Diabaté et al., 2005, 2008), and that a particular chromosomal arrangement in *An. coluzzii* performs well in low-rainfall environments (Touré et al., 1994). These conclusions, however, tend to be based on relatively small-scale observations or experiments. Some information on relative abundance at larger scales is available (della Torre et al., 2005; Caputo et al., 2008; Simard et al., 2009; Costantini et al., 2009), but very little modelling has been done to quantify these differences across sub-Saharan Africa. So far only occurrence information has been generally used (Sinka et al., 2010). In contrast, the relative abundance of *An. gambiae* s.s. in its former definition (including *An. coluzzii*) versus *An. arabiensis* has long since been modelled and estimated across sub-Saharan Africa (Lindsay et al., 1998).

A notable exception in this context is Tene Fossog et al. (2015), who develop a logistic regression model using abundance data of each subspecies across western sub-Saharan Africa to predict relative probability of occurrence of the two subspecies. They use model selection to select a subset of relevant spatial, climatic and land cover variables in their predictions. However, despite acknowledging the likelihood of nonlinear effects of some variables (see Figure 3b in their paper), they use only linear predictors in a logistic regression. We use their work as a starting point to develop a flexible neural network model to incorporate nonlinear relationships, along with additional predictors and newly collated records of relative abundance (VectorBase: Giraldo-Calderón et al., 2015) to extend our predictions to include the rest of sub-Saharan Africa.

### 1.4 Dispersal

Anopheline mosquitoes have historically been categorised as being unlikely to migrate long distances (with mean dispersal distances typically less than 1 km, maximum distances typically no greater than 5 km). Although longer range dispersal events are possible and have been linked to mosquito-borne disease outbreaks, short range dispersal is supposed to predominate life history strategies (Gillies, 1961; Service, 1997). A recent empirical study (Huestis et al., 2019), however, provides evidence for wind-driven long-range dispersal of *An. gambiae* s.s. and *An. coluzzii* mosquitoes in large numbers. These mosquitoes remain capable of reproduction and pathogen transmission (Sanogo et al., 2020), and are estimated to regularly travel much further than even a rare long-range dispersal event could achieve. Moreover, it has been recently suggested that *An. gambiae* s.l. populations in areas of low human density may also facilitate migration, gene flow or both (Epopa et al., 2020).

### 1.5 Aestivation

Another source of controversy is aestivation, in which mosquitoes become dormant during dry conditions in which they would not otherwise survive. While not proven to occur widely on a population scale, it is a popular hypothesis for wet season reemergence of *An. coluzzii* in the Sahel (Dao et al., 2014; Lehmann et al., 2017). Modelling studies that address this problem include Magombedze et al. (2018) and North and Godfray (2018). The latter simulation study notes that rare persistent water sources provide a competing explanation for persistence through the dry season, as does long distance migration. In this model, these latter proposed processes are accommodated by spatially and temporally varying carrying capacities (Section 1.3) and dispersal behaviour (Section 1.4); aestivation is not explicitly modelled.

### 1.6 Spatial Scope

The spatial scope for this analysis includes all countries within the African region as defined by the United Nations geoscheme that are within the range of *Anopheles gambiae* s.l. (United Nations regions are listed here: https://unstats.un.org/unsd/methodology/m49/overview/). The spatial scope includes the range of *Anopheles gambiae* s.l. on the African continent and also island countries or territories of the African region where *Anopheles gambiae* s.l. is present, such as Madagascar, Mauritius, Comoros and São Tomé and Príncipe. *Anopheles gambiae* s.l. is also found in Cabo Verde (DePina et al., 2018).

## 2 Methods

### 2.1 Spatio-temporal mosquito demographic model

We represent each combination of age class (*a*), sex (*s*), genotype (*g*) and subspecies of mosquito (*m*) as a separate scalar field *X*_*a,s,g,m*_(*t*, **x**) in a Partial Differential Equation (PDE) model with time *t* and 2D location **x**. Mosquitoes are assumed to not persist in ocean regions, and we set a zero-population Dirichlet condition in those regions. We allow, however, for the possibility that mosquitoes can advect across the ocean to neighbouring islands for a maximum duration of one day.

We represent the model numerically by spatially discretising the mosquito population of sub-Saharan Africa into 5 km × 5 km grid cells using the Africa Albers Equal Area Conic projection (ESRI:102022), and temporally discretising in 1 day timesteps, using fourth-order Runge-Kutta to integrate between timesteps. This timestep was determined experimentally to be both numerically stable and accurate. Combined with the spatial resolution, the model can be run within a reasonable time frame, while still being suitable for modelling the relevant biological processes, with a mosquito home range being roughly one cell in size. Within a timestep, we model demographic processes, followed by diffusion (see Section 2.1.4), followed by advection (see Section 2.1.5). While our model is deterministic, we allow extinction to occur by making a continuous model correction: at the end of the demographic processes step in each timestep, any cell with less than one total mosquito in a subspecies is set to zero for all classes of that subspecies.

We define *N*_*subspecies*_ subspecies, such that each subspecies is given an integer number from *m* = 0 to *m* = *N*_*subspecies*_: in our main results, *N*_*subspecies*_ is set to 2 to represent *An. gambiae* s.s. and *An. coluzzii* as *m* = 0 and 1 respectively. We also discretise age into *N*_*age*_ age classes in our model: in our main results, we use *N*_*age*_ = 2 (one newborn and one adult class), but we describe the general model for *N*_*age*_ ≥ 1 here, as different values of *N*_*age*_ may be more relevant for different model applications. For *N*_*age*_ = 2 age classes, the constant transition rate *b* results in an exponential distribution of maturation times between larval classes. Some larvae will quickly mature, potentially making the population more resilient to intervals without rainfall. Although we focus on *N*_*age*_ = 2, Figure S2 shows that results for *N*_*age*_ = 6 closely match results for *N*_*age*_ = 2 in Figure S1, with resistance spreading slightly more slowly over time.

Ages vary from the “newborn” larvae age class *a* = 0 (into which all individuals are born) through to adults at *a* = *N*_*age*_ − 1, with [1, *N*_*age*_ − 2] representing intermediate larval stages where these exist (*N*_*age*_ > 2). We here describe the PDE separately for these three stage types. Note that in our model we only track those mosquito eggs that produce viable larvae, hence our newborn class consists of larvae instead of eggs. Our numerical model is written in Cython (the Python programming language with C extensions Behnel et al., 2011) and visualisations are performed in the R programming language (R Core Team, 2020).

#### 2.1.1 Adults

The PDE governing each scalar field representing adult populations 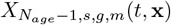 for *N*_*age*_ > 1 is given by

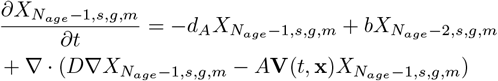

where subscript *s* denotes sex (*M* or *F*), *g* genotype, and *m* mosquito subspecies ((1) *An. gambiae* s.s. and (2) *An. coluzzii*). We model three alleles: *w* for wild–type, *c* for construct (gene drive system), and *r* for resistant. This results in a set of six potential genotypes *G* = {*ww, wc, wr, cc, cr, rr*}. The vector field **V**(*t*, **x**) represents the wind experienced across the spatial domain at a specified time *t* and place **x**. Note that our model makes some modifications to the advection process for biological reasons, specified below in Section 2.1.5. Other parameters are given in Table 1.

**Table 1:** Parameter definitions and estimates for final model (*N*_*age*_ = 2)

| Variable | Name | Description | Estimate | Source |
| --- | --- | --- | --- | --- |
| **Demographics** |  |  |  |  |
| $d_A$ | Adult mortality | Daily probability of mortality | $0.1 \text{ d}^{-1}$ | North and Godfray (2018) (or see Lunde et al. (2013), Eckhoff (2011), North et al. (2013), Wu et al. (2020)) |
| $d_J$ | Juvenile (larval) mortality | Daily probability of mortality | $0.05 \text{ d}^{-1}$ | North and Godfray (2018) (or see Lunde et al. (2013), Eckhoff (2011), North et al. (2013), Magombedze et al. (2018)) |
| $b$ | Larval transition rate | Number of days from an egg being laid to when it emerges as a sexually mature adult (when $N_{age} = 2$) or reaches next larval stage ($N_{age} > 2$) | $0.1 \text{ d}^{-1}$ | North and Godfray (2018) (or see Eckhoff (2011), North et al. (2013), Wu et al. (2020)) |
| $D$ | Diffusion coefficient | Typical rate of spread of population from a point source | $900 \text{ m}^2 \text{ d}^{-1}$ | Lunde et al. (2013) (or see Ickowicz et al. (2021)) |
| $\lambda$ | Larvae per female | Expected number of larvae per female per day (wild type mosquitoes) | $9 \text{ female}^{-1} \text{ d}^{-1}$ | North and Godfray (2018) (or see Lunde et al. (2013), Eckhoff (2011), Wu et al. (2020)) |
| $k$ | Relative probability of mating between subspecies | The relative probability that a female has offspring with a male of a different subspecies to her own ($k < 1$) | 0.01 | Pombi et al. (2017) (or see Lee et al. (2013)) |
| $\alpha_{ij}$ | Lotka-Volterra competition between subspecies | The relative effect on subspecies X of a member of a different subspecies Y taking up its resources (and thus larval carrying capacity), as compared to a conspecific | $\alpha_{11} = \alpha_{22} = 1$,<br>$\alpha_{12} = \alpha_{21} = 0.4$ | Pombi et al. (2017) |
| **Genetics** |  |  |  |  |
| $k_c$ | Probability of cleavage | | 0.995 | Beaghton et al. (2017) |
| $k_j$ | Probability of non-homologous repair | | 0.02 | Beaghton et al. (2017) |
| $k_n$ | Probability nuclease gene lost during homing | | $10^{-4}$ | Beaghton et al. (2017) |
| $h_n$ | Dominance coefficient for nuclease expression | | 0.5 | Beaghton et al. (2017) |
| $s_n$ | Cost of nuclease expression | | 0.05 | Beaghton et al. (2017) |
| **Larval carrying capacity** |  |  |  |  |
| $K_m(\mathbf{x})$ | Larval carrying capacity | | Details in text (2.2) and below parameters | |
| $\alpha_0(\mathbf{x})$ | Permanent larval site population | | 0 (no permanent sites) | North and Godfray (2018) |
| $\alpha_1$ | Contribution to breeding from rainfall | | 200,000 | North and Godfray (2018) |
| $\alpha_2$ | Larval sites associated with rivers and lakes | | 200,000 | North and Godfray (2018) |
| $\phi$ | Rate of carrying capacity population increase with rainfall | | 0.03 per mm rain per week | North and Godfray (2018) |
| $\kappa$ | Rate of carrying capacity population increase with water bodies | | 0.8 per km water | North and Godfray (2018) |
| $\delta$ | Replenishment rate of intermittent water sites with rainfall | | 0.03 per mm rain per week | North and Godfray (2018) |
| **Model details** |  |  |  |  |
| $X(0, \mathbf{x})$ | Initial condition | | Details in text (2.3.2) | |
| $A \mathbf{V}(t, \mathbf{x})$ | Advection | | Details in text (2.1.5, 2.3.1) | Huestis et al. (2019) |
| $t$ | Time domain for integration | | 2005-2015 | |

Here *N*_*age*_ − 1 indicates the adult age bracket. If *N*_*age*_ > 1 then *N*_*age*_ − 2 is the eldest larvae age bracket; the special case of *N*_*age*_ = 1 is considered below (Section 2.1.3). The first term on the right-hand side 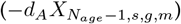 describes the mortality of adults, while the second term 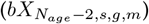 describes aging from the eldest larvae. The final term represents diffusion (∇*D*∇*X*) and advection 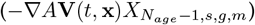, with *A* being the probability of an adult mosquito being advected by wind.

#### 2.1.2 Larvae

For *a* ∈ [1, *N*_*age*_ − 2], the populations are governed by

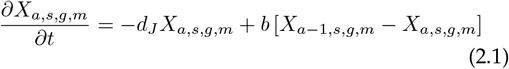

As above, the first term on the right-hand side (−*d*_*J*_*X*_*a,s,g,m*_) describes the mortality of this age-bracket of larvae, while the second (*b* [*X*_*a*−1,*s,g,m*_ − *X*_*a,s,g,m*_]) describes aging to/from older and younger age brackets respectively. Note that for *N*_*age*_ ≤ 2 there are no such intermediate larvae.

#### 2.1.3 Newborns and N_age_ = 1 model

The PDE governing *X*_0,*s,g,m*_(*t*, **x**) is given by

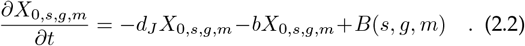

Where *N*_*age*_ = 1, there is no age structure and the full PDE takes the form

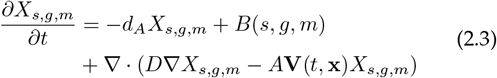

Note that for readability, we only use subscripts when referring to mosquito population classes (i.e. *X*_*a,s,g,m*_) and refer to parameters which differ by population class as functions of the offspring classes. Note that we do not include parental classes (i.e. *g*_*M*_, *g*_*F*_, *m*_*M*_, *m*_*F*_) in the function names, also for brevity, and that these are only defined explicitly in the function definitions.

In Equation 2.2, the first term (−*d*_*J*_*X*_0,*s,g,m*_) describes mortality of newborns, while the second (−*bX*_0,*s,g,m*_) describes aging into the youngest age-bracket of larvae (or adults for *N*_*age*_ = 2). The final term describes the birth of newborn larvae of a given sex *s*, genotype *g* and subspecies *m*, given all possibilities of the mother’s and father’s genotype and subspecies (*g*_*M*_, *g*_*F*_, *m*_*M*_ and *m*_*F*_; described in more detail later). For brevity, we describe it as the product of functions describing the relevant biological mechanisms:

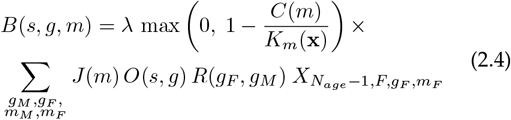

where the baseline fecundity rate is *λ*, the expected number of larvae per clutch of eggs per female per day, assumed produced by a mating of wildtype mosquitoes. The *J* and *O* terms represent subspecies inheritance and genotype inheritance (including sex) respectively, and are described below. In our main results, we keep the relative fecundity function *R*(*g*_*M*_, *g*_*F*_) constant at *R* = 1, but this can readily be varied to represent reduced fecundity due to inviability of a gene drive construct — see Supplementary Materials Figure S3 for an example.

The max 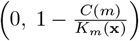 term models the effect of *K*_*m*_(**x**), the (spatially varying) larval carrying capacity for subspecies *m*, on the fecundity of that subspecies using a logistic function. The inclusion of a max term is to keep the model biologically plausible: without it, *C*(*m*) > *K*_*m*_(**x**) would mean that negative newborns are produced. Here *C*(*m*) is the competition that a newborn of subspecies *m* experiences from all larval populations:

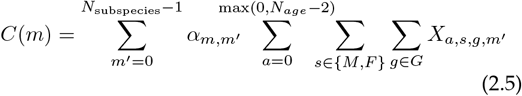

Where 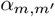 represents the effect of competition of subspecies *m* on subspecies *m*′, and the max term is here used to ensure that where *N*_*age*_ = 1, the larval carrying capacity instead applies to the adult population.

The function *J* (*m*) returns the probability that a male adult of genotype *g*_*M*_ and subspecies *m*_*M*_ successfully mates with a female adult of genotype *g*_*F*_ and subspecies *m*_*F*_ to produce newborns of subspecies *m*:

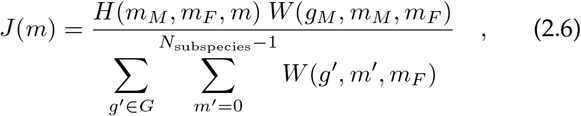

where the number of available males of a subspecies and genotype is

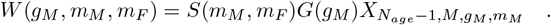

The numerator is the relative number of expected matings between a male of subspecies *m*_*M*_ (through *S*) and genotype *g*_*M*_ (through *G*) and the given female, while the denominator normalises the probability.

The relative probability of mating based on subspecies *S*(*m*_*M*_, *m*_*F*_) = 1 if *m*_*M*_ = *m*_*F*_ and *k* otherwise. The relative fitness based on genotype is:

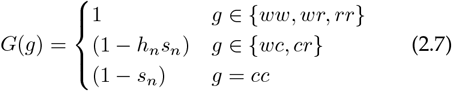

where *h*_*n*_ and *s*_*n*_ are adapted from the Beaghton et al. (2017) model (see Table 1).

We compare two different scenarios for subspecies inheritance *H*(*m*_*M*_, *m*_*F*_, *m*); maternal inheritance and equal inheritance. For maternal inheritance, the proportion of offspring born of a mating between subspecies *H*(*m*_*M*_, *m*_*F*_, *m*) = 1 if *m*_*F*_ = *m* and 0 if *m*_*F*_ ≠ *m* (mother is always the same subspecies as her offspring). For equal inheritance, the proportion of offspring born of a mating between subspecies *H*(*m*_*M*_, *m*_*F*_, *m*) = 0.5 if *m*_*M*_ ≠ *m*_*F*_ (parents are different subspecies, i.e. cross-species offspring are equally split between subspecies). In both scenarios, *H*(*m*_*M*_, *m*_*F*_, *m*) = 1 if *m*_*M*_ = *m*_*F*_ = *m* (both parents and offspring same subspecies) and 0 if *m*_*M*_ ≠ *m* and *m*_*F*_ ≠ *m* (both parents different subspecies to offspring). Where there are more than two subspecies, we also need to specify that *H* = 0 when *m*_*M*_ ≠ *m*_*F*_, *m*_*F*_ ≠ *m* and *m* ≠ *m*_*M*_ (i.e. parents and offspring are all of different subspecies).

The function *O*(*s, g*) gives the probability of sex *s* and genotype *g* for the offspring. This probability depends on the genotypes of the parents such that

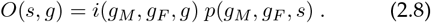

The first term *i*(·) describes the probability of an offspring inheriting genotype *g* given parents of genotypes *g*_*M*_ and *g*_*F*_. We again adapt the Beaghton et al. (2017) model to our target genotypes, including the effect of the gene drive construct, and using their parameter values (see Table 1). However, instead of assuming full random mixing of alleles as in their model, we directly model genotypes of each parent (see Table 2).

**Table 2:**
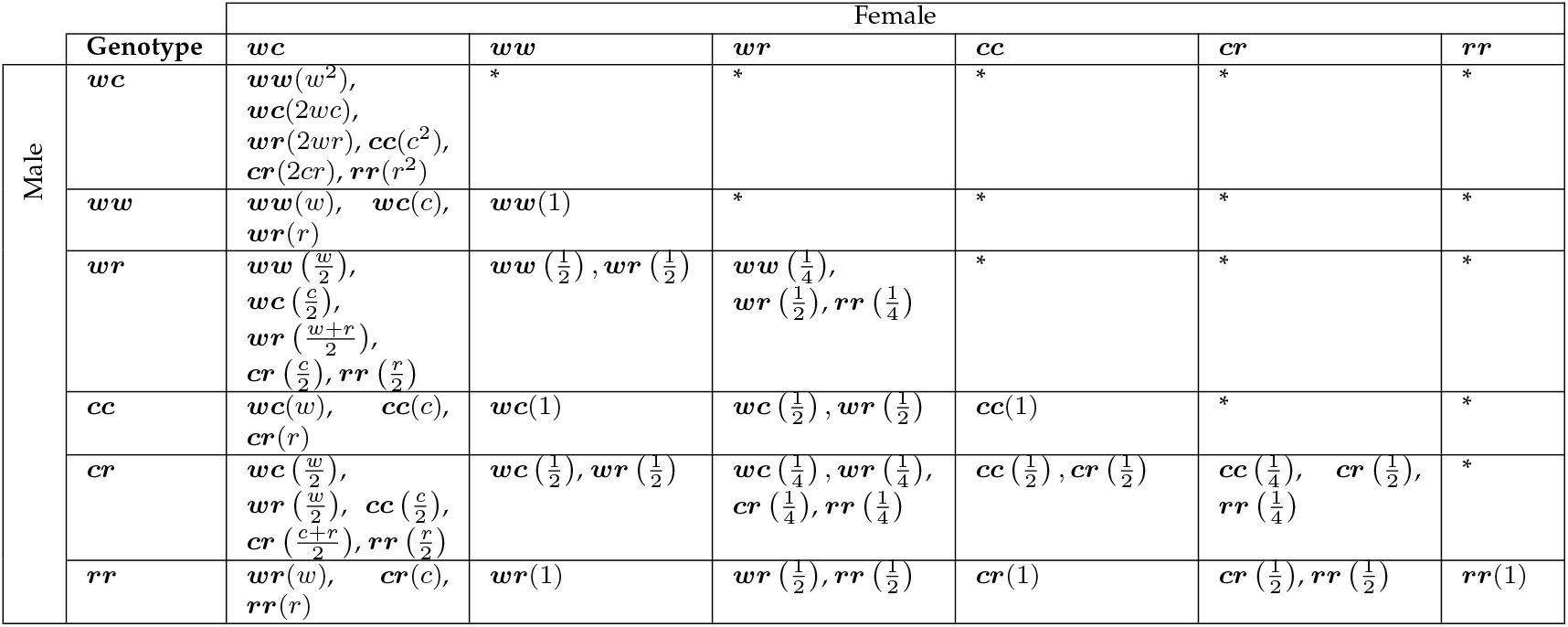
Possible values for each offspring *g*′ in inheritance table *i*(*g*_*M*_, *g*_*F*_, *g*′), with probability of occurrence given by the number in parentheses, where allele probabilities *w* = 1*/*2 *− k*_*c*_*/*2, *c* = 1*/*2 + *k*_*c*_(1 *− k*_*j*_)(1 *− k*_*n*_)*/*2 and *r* = *k*_*c*_(*k*_*n*_ + *k*_*j*_ (1 *− k*_*n*_))*/*2. Table is symmetric, so cells marked with * have the same values as their transposes.

The second term *p*(·) describes the proportion of male offspring (and thus sex bias mechanisms). In the results presented in the main paper, we keep *p* constant at *p* = 0.5, though this can be readily varied to model constructs that induce sex bias in viable offspring (see Supplementary Materials Figure S4 for an example). The sex ratio is kept constant between subspecies (Paaijmans et al., 2009).

#### 2.1.4 Diffusion

Diffusion is modelled by using the nearest-neighbour finite-difference approximation to the Laplacian. That is, with timestep size Δ*t* and cell length Δ*x*, the fraction of diffusing mosquitoes removed from one cell is 4*D*Δ*t/*(Δ*x*)^2^, where Δ*x* is the cell-size. One quarter of this amount is added to each of the 4 neighbours. If the neighbours happen to be inactive (such as on the model boundary) those mosquitoes are assumed to die instantly.

#### 2.1.5 Wind advection

Wind advection is expected to occur over very short time periods (overnight, as mosquitoes are not believed to travel during daylight; see Huestis et al., 2019) and potentially very large distances (hundreds of kilometres, passively carried by the wind with negligible resistance; see Huestis et al., 2019). As such, we explicitly trace the trajectories of mosquitoes from each cell during each timestep (1 day), as advected by the wind vector field **V**(*t*, **x**). We use the Cross-Calibrated Multi-Platform (CCMP) Ocean Surface Wind Vector Analyses dataset (Atlas et al., 2011) to define this vector field; the data is available at the required daily timesteps over the time period required. We interpolate their vector field, given at 0.25 degree intervals (approximately 28 km at the equator), to fit our grid. Huestis et al. (2019) set up two different scenarios in their own modelling, where mosquitoes mosquitoes travel either 2 hours or 9 hours a night. We explore both scenarios here, and also a third scenario with no wind advection.

### 2.2 Larval carrying capacity

#### 2.2.1 Total abundance

We apply the method of North and Godfray (2018) to estimate larval carrying capacity for both species combined. Specifically, we use Equation 1 from their paper:

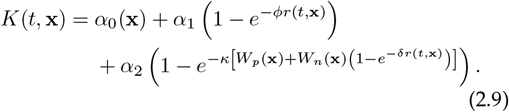

with parameters as estimated in their paper using Markov Chain Monte Carlo simulation with population data (see Table 1). The model incorporates rainfall (*r*(*t*, **x**) in mm per week), as well as the availability of nearby rivers and lakes; these may be either permanent or intermittent (*W*_*p*_(**x**) and *W*_*n*_(**x**) respectively; details in their paper). It also has scope for modelling permanent larval sites (*α*_0_(**x**)) but we set this to be zero for our model. Whereas North and Godfray (2018) uses settlements as their sites for mosquito populations, we are calculating populations across a square grid, so we assume that each cell contains a settlement as modelled in their paper if and only if there is a human population present in the cell. An implicit assumption in this approach is that areas without human settlements yield zero carrying capacity. This assumption is explored for a sparsely populated region as follows. To calculate *K*(*t*, **x**) at each cell at each required timestep using Equation 2.9, we estimate human presence or absence by using data produced by the Facebook Connectivity Lab and Center for International Earth Science Information Network - CIESIN - Columbia University (2016) and publicly available from the Humanitarian Data Exchange (HDX; https://data.humdata.org/), except in Sudan, South Sudan and Somalia where these data are not available. There we assume human presence in all cells, which as an extreme case would facilitate the spatial spread of populations by local dispersal or advection. The other extreme, assuming human absence, is explored in Supplementary Materials Figure S7: it was found to have minimal effect outside of these countries, which take up a relatively small area near the edges of the species range.

We use inland water data from the Digital Chart of the World (DCW) as in their paper, and obtain monthly rainfall data from NASA’s Land Data Assimilation System (https://ldas.gsfc.nasa.gov/fldas). We also set the mosquito population and carrying capacity to zero for each subspecies in locations outside their range as estimated by the Malaria Atlas Project (see Wiebe et al., 2017, updated by the Malaria Atlas Project, accessed 25 May 2021 from https://malariaatlas.org/ and available for use under a Creative Commons Attribution 3.0 Unported License, https://creativecommons.org/licenses/by/3.0/legalcode).

#### 2.2.2 Relative abundance

Once we have a measure of the combined larval carrying capacity *K*(*t*, **x**), we need to separate this for two subspecies *K*_1_(*t*, **x**) and *K*_2_(*t*, **x**), which requires knowledge of the relative abundance of the two subspecies at each site. We use available field data collated by Tene Fossog et al. (2015) on the number of captures of both subspecies at various locations across Africa to flexibly predict a spatially-varying but temporally-constant relative abundance. We call this relative abundance *K*_*r*_(**x**) = *K*_1_(*t*, **x**)*/K*(*t*, **x**), or the proportion of *An. gambiae* s.s. at a site.

We first attempt to replicate the results of the Tene Fossog et al. (2015) logistic regression model by independently sourcing the predictors that they used in their final model: these are Latitude, Distance to Coast, Annual Mean Temperature, Mean Temperature of Wettest Quarter, Mean Elevation, Annual Normalized Vegetation Difference Index (NVDI) and Annual Variation in NVDI. We then apply a logistic regression model to the (slightly different) predictors, as they did. As well as providing an independent verification of the results in their paper, sourcing the predictors ourselves allows us to use more data sources across sub-Saharan Africa, allowing us to extrapolate and test different modelling approaches.

Using our independently sourced version of the predictors, we then apply a dense feed-forward neural network to flexibly model the relative abundance function *K*_*r*_(*x*) (see Supplementary Materials Appendix S2 for details). We perform leave-one-out cross-validation (jackknifing) on each of these models, comparing the replicated logistic regression model and the original Tene Fossog et al. (2015) model with the neural network model, to test whether there is an increase in performance by incorporating nonlinear effects on the same set of variables.

The Tene Fossog et al. (2015) model uses only allopatric sites (those where only one subspecies was detected) to train their models, as did our replication model and initial neural network model. As we are interested specifically in subspecies overlap, we include sympatric sites (where both subspecies were detected), including three new sites (Antonio-Nkondjio et al., 2012, 2013) publicly available from VectorBase (Giraldo-Calderón et al., 2015). We also add further predictors of potential interest to the described neural network model: mean annual precipitation (BIO12) and precipitation of wettest quarter (BIO16), which were also of interest to their model but excluded by their model selection process, along with salinity (Pombi et al., 2017) which was discussed in detail by Tene Fossog et al. (2015) but not included as a model predictor. We then apply forward selection to select the model variables in the final model (see Supplementary Materials Appendix S2 for details). To ensure convergence in the larger range of scenarios, we here use a 50:50 training-testing split on the data, but otherwise keep the same network topology and approach. Once this process is complete, we then have our final estimate of *K*_*r*_(**x**), which we can then apply to the previously calculated *K*(*t*, **x**) to obtain results for both *K*_1_(*t*, **x**) and *K*_2_(*t*, **x**).

### 2.3 Parameters

Where possible and reasonable, we take parameter estimates from literature for our model (see Table 1), while checking that multiple sources give similar results.

#### 2.3.1 Wind advection

We use estimates from Huestis et al. (2019) to indirectly estimate the probability of adult mosquitoes being advected by wind. They estimate that 6 million *An. coluzzii* mosquitoes cross a 100 km line perpendicular to the prevailing wind direction every year. Over the course of a 2 or 9 hour flight (the two night-time flight times tested in their Methods) their calculated trajectories give displacements of 3–69 km and 47–270 km respectively (means 38.6 and 154.1; 95% mean CIs 37–41 and 140– 168). If we assume that each migrating individual completes just one 2 or 9 hour overnight flight each way and are only counted once, the mosquitoes that will cross the imaginary 100 km line will largely come from a rectangle bounded by this line on one side, and another 100 km line positioned either 38.6 or 154.1 km upwind on the other side. So the area over which mosquitoes will cross this line will be roughly 100 km × 38.6 km = 3, 860 km^2^ or 100 km × 154.1 km = 15, 410 km^2^, within which 6 million *An. coluzzii* migrating mosquitoes are estimated to cross. This gives us a density of 1.55 × 10^− 3^ or 3.89 × 10^− 4^ migrating mosquitoes per square metre per year: converting to relevant units gives us 106.39 and 26.65 mosquitoes per 5 km × 5 km cell per daily timestep. If we can then estimate the overall density of mosquitoes at their capture sites, we can then estimate the daily migration rate. Using the total abundance model from North and Godfray (2018), the average carrying capacity for larvae at coordinates 14°N, 6.7°W (located in between the capture sites in Mali) across the simulation period (2005–2015) is 41, 526. Given the other parameters *d*_*J*_ = 0.05 d^− 1^, *d*_*A*_ = 0.1 d^− 1^ and *b* = 0.1 d^− 1^, at equilibrium we would expect approximately the same number of adults as larvae at carrying capacity. This gives a daily migration rate of 106.39*/*41, 526 ≈ 2.6 × 10^− 3^ (or 1 in 390) for 2 hour migration; and 26.65*/*41, 526 ≈ 6.4 × 10^− 4^ (or 1 in 1558) for 9 hour migration.

#### 2.3.2 Initial conditions

As we expect similar numbers of larvae and adults, we set the initial conditions of the model for each age *a*, sex *s*, genotype *g* and mosquito subspecies *m* to

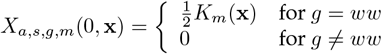

In cells where both subspecies exist, there will initially be competitive effects reducing numbers of one or both subspecies. In addition, advection and diffusion will affect the equilibrium. For these reasons, we run the model once from the initial condition described for the 11-year time period from 1 January 2005 to 31 December 2015 as a “burn-in” period, in order for the population to approximately reach a dynamic equilibrium that might be encountered in the wild.

Once the burn-in period is complete, we simultaneously introduce 10,000 male mosquitoes of each subspecies (*An. gambiae* s.s. and *An. coluzzii* that are heterozygous with the construct (genotype *wc*) in 15 separate locations (see Figure 4 in the Results), and run the model for another 11-year period. We choose heterozygous mosquitoes to introduce as these have less fitness cost than homozygous mosquitoes (see Equation 2.7), increasing the chance of spread. The first five of these are placed on islands off the African mainland at the nearest suitable location (see below) to assess the potential for incursion of the genetic construct onto the mainland. The island sites chosen as illustrative examples are:

1. the Bijagós islands (off Guinea-Bissau),
2. Bioko (off Cameroon),
3. Zanzibar (off Tanzania),
4. Comoros (off Mozambique), and
5. Madagascar.

The other ten sites (numbered 6–15) were chosen to be as evenly spaced as possible across the range of *An. Gambiae* s.s. where at least 10,000 mosquitoes of either subspecies is available year-round (the algorithm is described in Supplementary Materials Appendix S1).

### 2.4 Scenario tests

As mentioned in Section 2.1.5, we use the two scenarios from Huestis et al. (2019) that mosquitoes are passively advected with the wind for either 2 hours or 9 hours a night. We also add a third scenario of no wind advection at all, to fully explore the effect of wind on mosquito movement. These three scenarios are then combined with the two scenarios of subspecies inheritance (maternal and fifty-fifty) described in Equation 2.6 to make six total scenarios modelled.

## 3 Results

### 3.1 Larval carrying capacity

#### 3.1.1 Relative abundance

To compare the effectiveness of the modelling approaches on the jackknifed mosquito data from Tene Fossog et al. (2015) and VectorBase (Antonio-Nkondjio et al., 2012, 2013; Giraldo-Calderón et al., 2015), where the model probability of the subspecies being *An. coluzzii* at site *i* of *N* sites is *p*_*i*_, and the true probability is *s*_*i*_ (either 0 or 1 depending on subspecies present), we use four measures:

- the actual misclassification rate or “error rate”, where the subspecies at site *i* is predicted to be *An. coluzzii* if *p*_*i*_ > 0.5, otherwise *An. gambiae* s.s.;
- the cross-entropy, calculated as

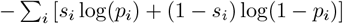
- the expected number of misclassifications or errors,

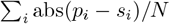
- and the root mean square (RMS) error,

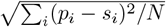

Table 3 shows that our neural network model performs as well (error rate), slightly better (expected error rate and RMS error) or much better (cross-entropy) on all of the measures. Its much better performance on cross-entropy is likely due to the fact that it is trained specifically to minimise cross-entropy, which may not directly correlate with lower error rates — for example, compared to the other measures, cross-entropy will much more harshly penalise a model for assigning a very low probability to an event which then occurs in the testing data.

**Table 3:** Comparison of modelling approaches using four different measures of accuracy. The “Original” model is that of Tene Fossog et al. (2015), the “Replication” is our attempt to replicate their model with available data for predictors, and “NN” is our neural network model as described in this paper, but using their predictors. The models are compared with the dataset described in their paper (Tene Fossog et al., 2015) and from VectorBase (Antonio-Nkondjio et al., 2012, 2013; Giraldo-Calderón et al., 2015))

|  | Original | Replication | NN |
| --- | --- | --- | --- |
| Error rate | 0.1051 | 0.1018 | 0.1051 |
| Cross-entropy | 170.06 | 172.43 | 158.30 |
| Expected error rate | 0.1595 | 0.1610 | 0.1513 |
| RMS error | 0.2839 | 0.2839 | 0.2728 |

Figure 1 illustrates the forward selection process. The chosen variables are Mean Annual Temperature, Latitude, Elevation and Distance to Coast — after this point, even the best chosen variable added to the model only increases the mean validation loss.

**Figure 1:**
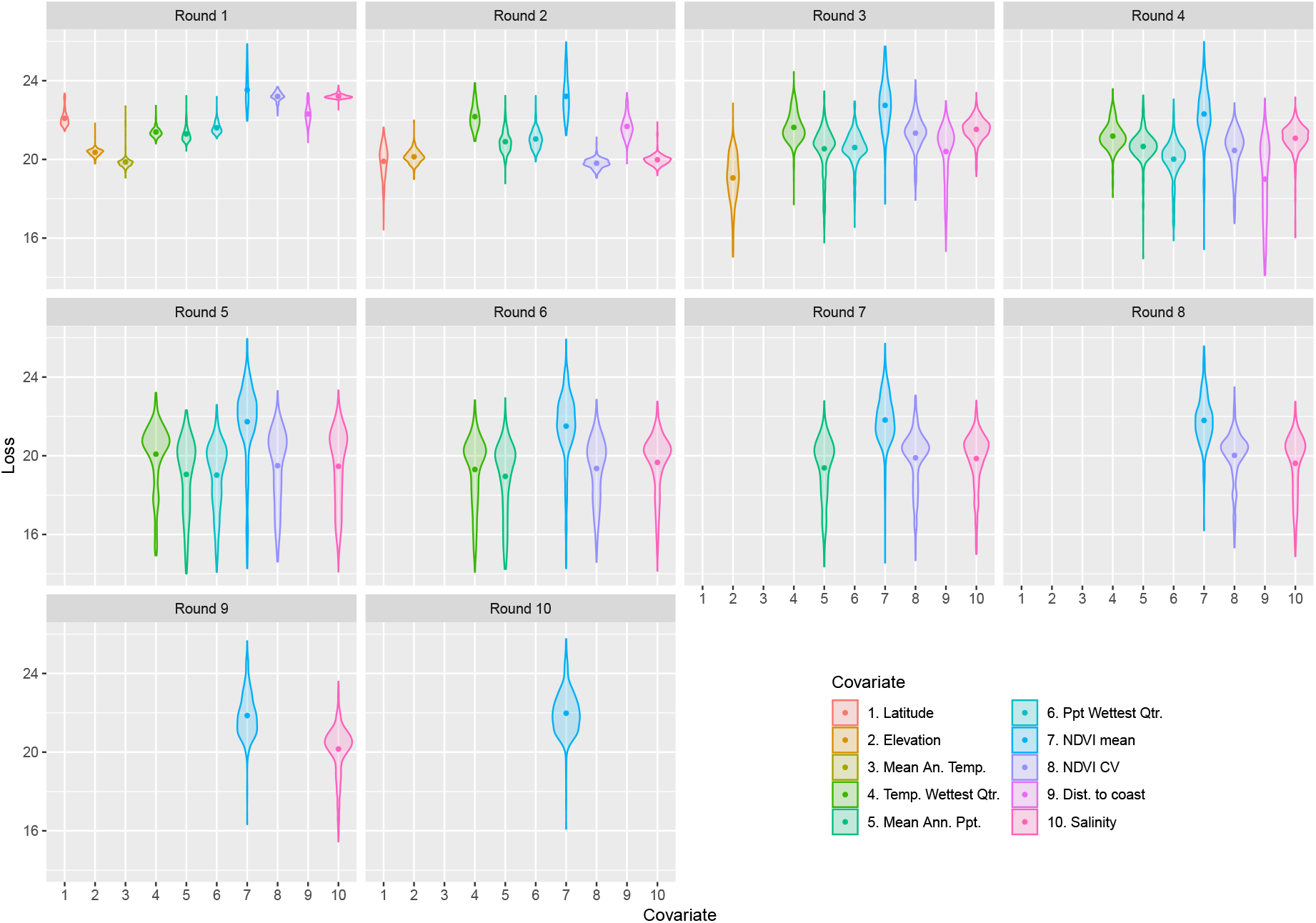
The forward selection process for the neural network model that estimates the relative abundance of the two subspecies. For each round of selection, the validation loss for each variable is shown, both for the 500 individual neural network runs (small dots) and the mean (large labelled dot), with a different colour. After a given round, the variable with the lowest mean validation loss is selected then the process is repeated with a model including that variable, testing all remaining variables. 95% bootstrap confidence intervals for the mean are displayed for each variable in each round.

Note that the NVDI Coefficient of Variation (CV) actually had the lowest mean validation loss in the second round, but Latitude was instead selected for four reasons:

- The two had mean validation losses that were statistically indistinguishable even after 500 model runs, based on the bootstrap 95% confidence intervals,
- Comparing the results of the forward selection process where either the NVDI CV or Latitude is chosen in the second round, the latter gives much better mean validation loss results in *subsequent* rounds, demonstrating that the forward selection process is sub-optimally choosing the NVDI CV,
- Data for the NVDI CV is much harder to source, and the variable is a less directly biologically relevant predictor than Latitude, and
- The NVDI CV becomes a much less desirable predictor after Latitude is selected, suggesting that it has little to contribute to the model that is unique to it, and not already contributed by Latitude.

Figure 2 shows the final estimated relative abundance as a mean of 100 neural network model runs using the chosen variables, alongside the standard deviation of these runs. The relative abundance follows a similar pattern to that of Tene Fossog et al. (2015), with *An. gambiae* s.s. mostly dominant except in some coastal areas and towards the Sahel. As expected, the variance between model runs is mostly low around the data points in western sub-Saharan Africa upon which the model was trained, with much higher variance around central and southern Africa. Interestingly the model runs also yield low variance around Ethiopia in particular, where they predict *An. gambiae* s.s. to be dominant.

**Figure 2:**
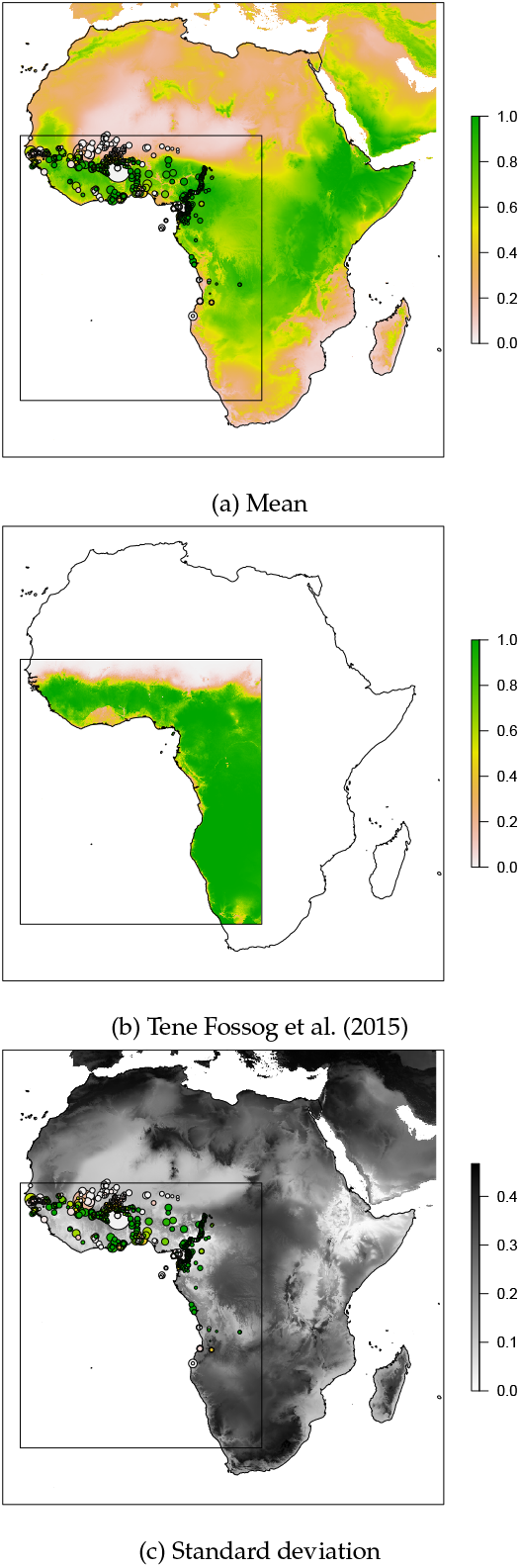
Summary statistics of relative abundance based on 100 neural network model runs. Mean relative abundance (a) is given as the proportion of *An. gambiae* s.s. (as opposed to *An. coluzzii*) in the local mosquito population, with low proportions given as white and high proportions as green. Circles represent data points on which the model is trained, filled with colour representing the proportion measured at the given site. The corresponding results from Tene Fossog et al. (2015) are given for the purposes of direct comparison (b). The standard deviation of relative abundance (c) is shown in greyscale, with white as low and black as high variance between model estimates at a given site. The rectangle denotes the area of study used in Tene Fossog et al. (2015).

#### 3.1.2 Total abundance

Figure 3 demonstrates the resulting estimated larval carrying capacity for each subspecies across the first year of modelling given the relative abundance model above and the North and Godfray (2018) total abundance model, combined with information on human presence and the known distributions of each subspecies.

**Figure 3:**
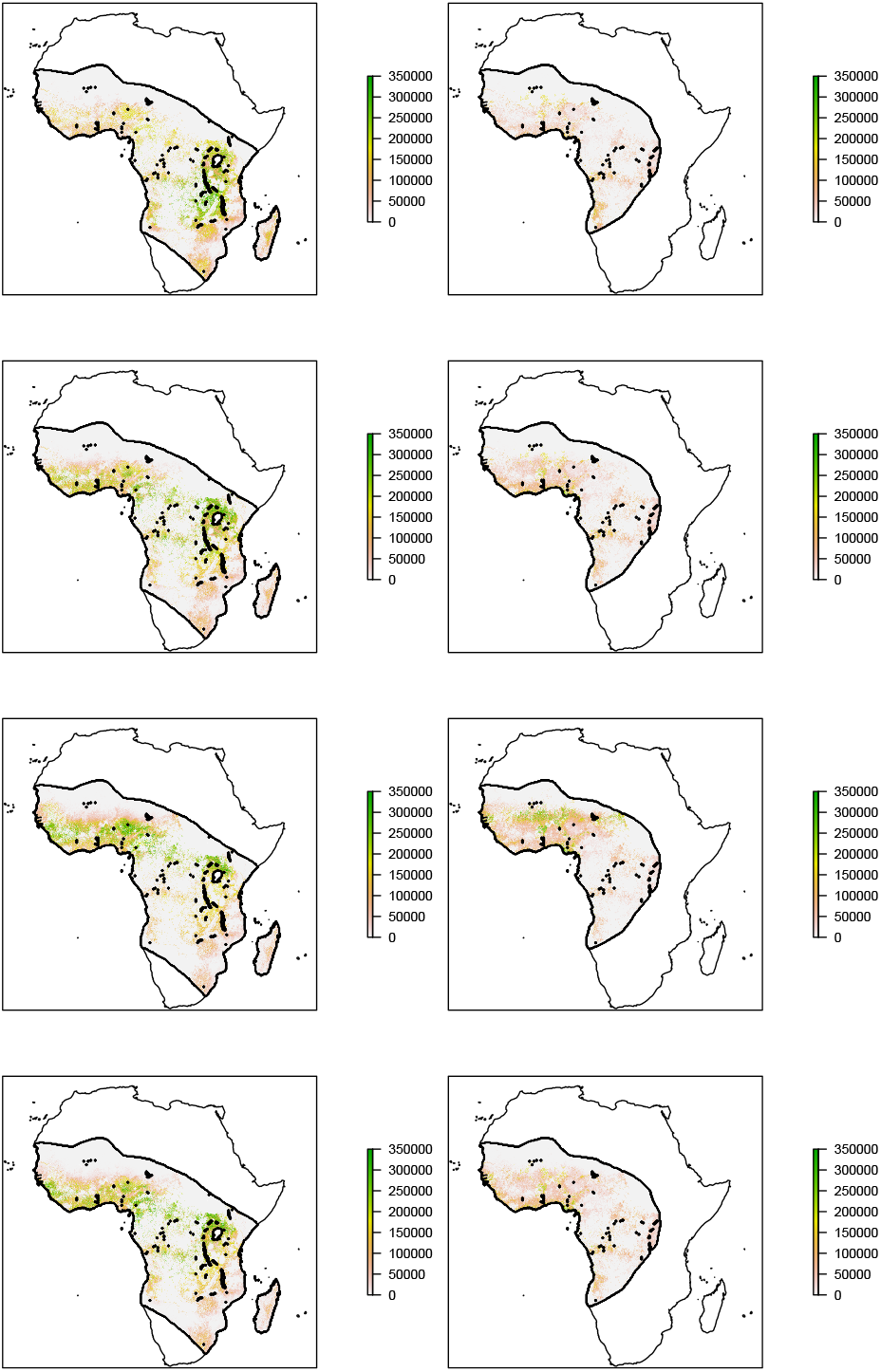
Estimated larval carrying capacity of *An. gambiae* s.s. (left) and *An. coluzzii* (right), for 2005 (the first year of modelling) in January (Southern Hemisphere summer), April (autumn), July (winter) and October (spring) from top to bottom.

### 3.2 Scenario tests

The choice of subspecies inheritance made no noticeable difference to the results for any of the three wind scenarios. Results shown use the maternal inheritance model.

Within the three wind scenarios, the zero-advection scenario experiences no noticeable mosquito transport on a continental scale, demonstrating that with the current selected parameters, advection is far more important than diffusion. We thus show figures for the 9-hour case below, contrast the 2-hour results in text, and present the full 2-hour and zero-advection results in Supplementary Materials.

Figure 4 shows the location and spread of the construct from the 15 introduction sites of the genetic construct (labelled 1–15) for the highest wind advection scenario (9 hours).

**Figure 4:**
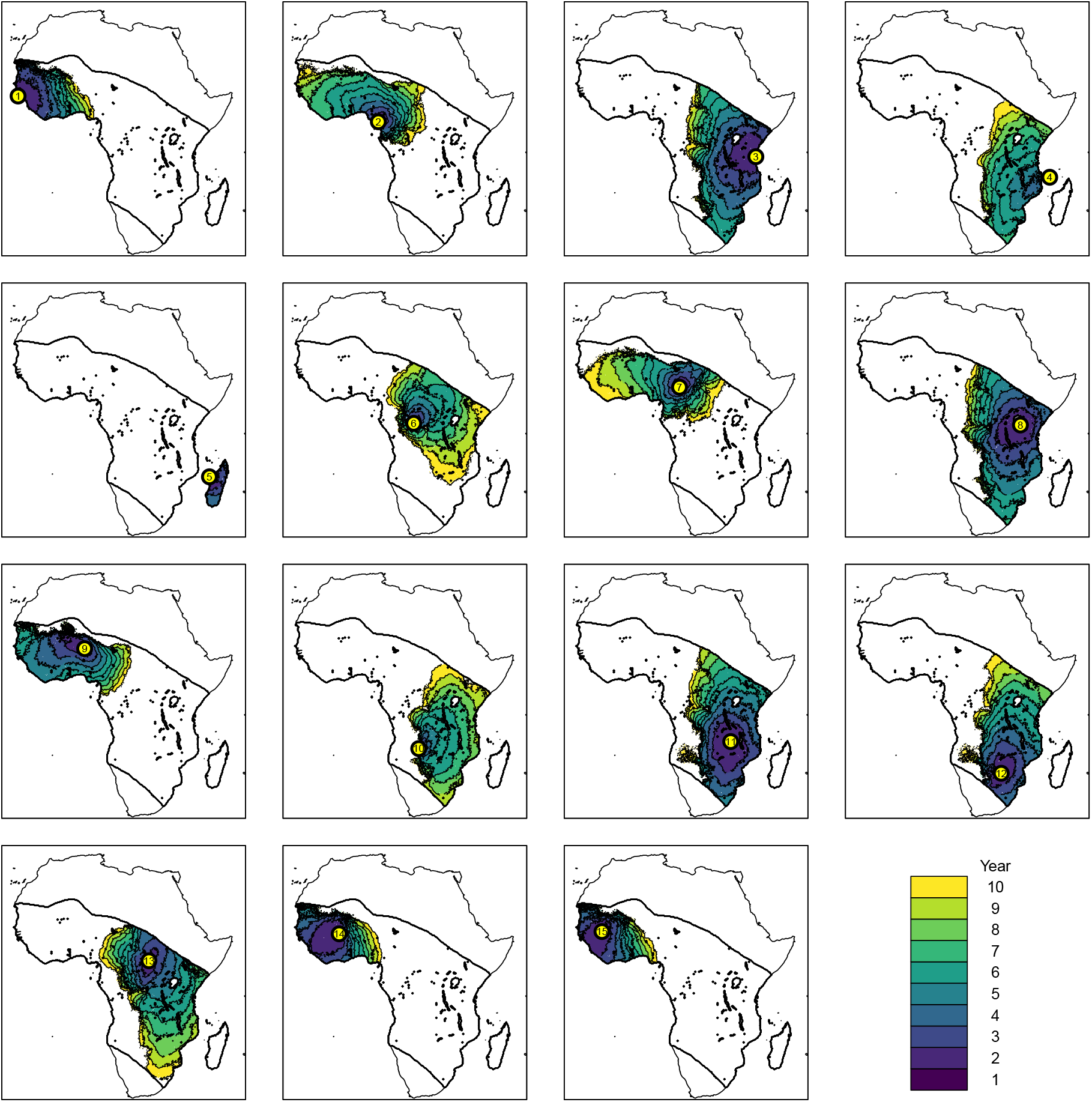
The invasion front of the construct (defined as having at least two alleles, e.g. one *cc* or two *wc* mosquitoes, in a cell) from a selection of starting points, with a separate colour given for each year. The island introductions are (1) the Bijagós islands (off Guinea-Bissau), (2) Bioko (off Cameroon), (3) Zanzibar (off Tanzania), (4) Comoros (off Mozambique) and (5) Madagascar.

All of the islands were able to maintain a population with the genetic construct (Figure 5), with all except for Madagascar (Site 5) then invading the African mainland in subsequent years. There appears to be a barrier to dispersal in central Africa, with most introductions either remaining in western Africa (sites 1, 2, 7, 9, 14 and 15) or eastern Africa (sites 3, 4, 6, 8, 10–13). When advection was reduced to 2 hours, only the Bijagós (Site 1) and Zanzibar (Site 3) introductions were able to reach the mainland (see Supplementary Materials Figure S6).

**Figure 5:**
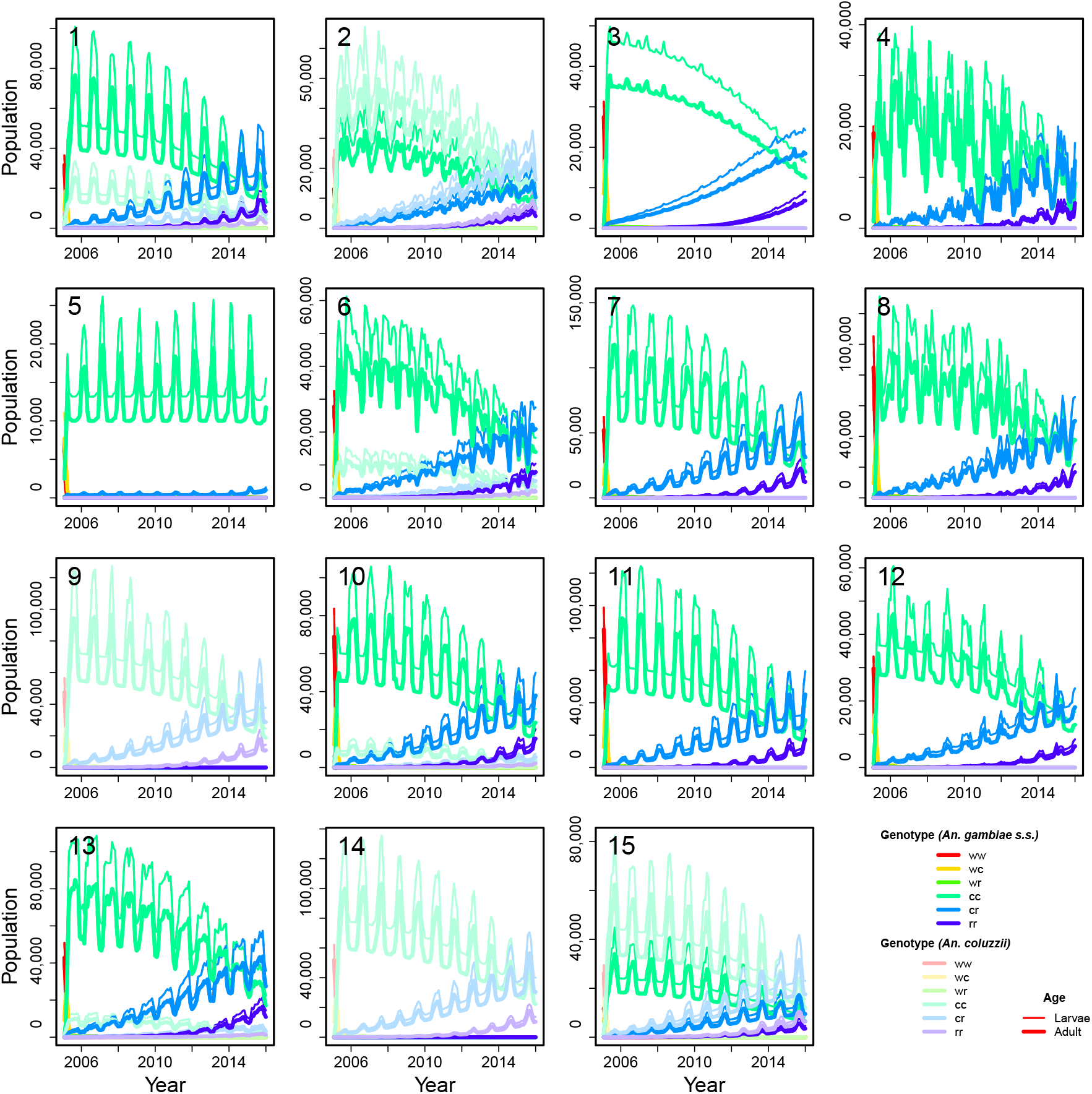
The time series abundance of male mosquitoes at each introduction point (the sub-figure number corresponds to the release site; see Figure 4), separated by species, genotype and age (female mosquitoes occur in identical numbers to males in this model). The colours correspond to genotype and the line thickness to age class.

Figure 5 shows more detail about the introductions at each site. At all sites, the construct allele completely takes over from the original wildtype allele in a matter of months. Once the construct is established in the population, resistance builds slowly but surely: the heterozygous resistant genotype *cr* becomes noticeable after a couple of years, and is beginning to overtake the wildtype by the end of the 11-year simulation period, with the homozygous resistant genotype (*rr*) starting to become noticeable in the population. All sites show a similar pattern, despite differences in scale, subspecies composition and seasonality. Only *An. gambiae* s.s. persists in Sites 3, 4, 5, 7, 8, 11 and 12, and conversely only *An. coluzzii* persists in Sites 9 and 14. The two coexist in the remaining sites (1, 2, 6, 10, 13 and 15). All sites show a regular seasonal pattern to abundance based on rainfall (the only seasonal driver in the model, other than wind), with the possible exceptions of Sites 4, 6 and 8, which show slightly more variable seasonality patterns. Why this might be is unclear, though these sites are all in central to eastern Africa.

Full colour animations of all process model outputs are available in Supplementary Materials.

## 4 Discussion

This study is the first continental scale model of population dynamics for two of the dominant malaria vector species in the *Anopheles gambiae* sensu lato species complex. The transient spread and persistence of a population replacement gene drive was predicted for the two hybridising subspecies *An. gambiae* sensu stricto and *An. coluzzii*. The two major factors that determine the spread of the gene drive at the continental scale were 1) potential wind-driven dispersal and 2) spatially and temporally varying carrying capacities for the two subspecies with both intraspecific and interspecific density dependence occurring at the larval stage. Our presented results are intended to demonstrate the plausibility of wide-scale spread of gene drive and structurally evaluate hypotheses of relevant life history strategies for these two dominant malaria vectors. The spatially explicit process model is designed to support more specific scenario-based assessments of both genetic and conventional vector control strategies.

Vertical gene transfer among the investigated subspecies is complicated by the spatially heterogeneous population structure, where introgression rates between *An. gambiae* s.s. and *An. coluzzii* vary with geographic location (*Anopheles gambiae* 1000 Genomes Consortium, 2017; Pombi et al., 2017; Clarkson et al., 2020), and perhaps also over time (Lee et al., 2013). Pombi et al. (2017) suggests that the rate of hybridisation between these subspecies depends on their relative frequency in the population. The model therefore tracks the relative number of available males by subspecies, and two alternative choices for species assignment of the first generation hybrids was investigated. The model results were found to be robust to species assignment via maternal descent when compared to equal proportions of species among the first generation hybrids.

The home range of *An. gambiae* s.l. is typically believed to be less than 5 square kilometres, although there is evidence for long-range dispersal (Huestis et al., 2019). Spatial dispersal was modelled through two different mechanisms: a local diffusive process, which may be mediated by the presence of *An. gambiae* s.l. between human settlements (Epopa et al., 2020), versus wind-driven advection. More generally, wind-driven dispersal events have been observed for mosquitoes including over ocean, although it is not understood whether or not wind-driven dispersal is a deliberate life history strategy for some species (Service, 1997). For genetic vector control strategies, the possibility of long range dispersal events are an important consideration that should be incorporated into the selection of field sites (James et al., 2018; Lanzaro et al., 2021). Unsurprisingly our results indicate that passive wind-driven advection, if present, can greatly increase the speed and spatial footprint of the invasion front for a near-neutral population replacement gene drive (Figure 4, Section 3.2).

The importance of wind-driven advection as a dispersal strategy for *An. gambiae* s.l. will likely be a key uncertainty in future risk assessments for genetic control strategies. The large difference in the dispersal between the 9 hour (Figure 4) and 2 hour results (Supplementary Materials Figures S5 and S6) have important implications for regional (trans-national) governance arrangements, the scope of stakeholder engagement activities and the degree of geographic containment that islands might provide during a staged-release strategy. In our simulated releases of the gene drive on islands, only Madagascar was sufficiently distant to prevent spread to the mainland under the 9 hour wind-assisted dispersal. Even when reducing wind-assisted dispersal to 2 hours, two of the five island releases resulted in spread to the mainland. Simulated introductions onto islands such as Bioko (32 km from the mainland) spread to the mainland, but only under the 9 hour scenario. Hence, only this result is consistent with recent genomic analysis that shows mosquito populations on this island are not isolated from the mainland (Campos et al., 2021). Although our results make the potential scale of wind-mediated spread of gene drive clear, exactly how and where this spread occurs in the relevant mosquito taxa, and the mosquito behaviour that helps drive it, is still being studied (Huestis et al., 2019; Florio et al., 2020). In addition, while data on previous wind patterns is readily available, predicting future wind patterns can only be done in very general terms, especially given the added complications and uncertainties caused by anthropogenic climate change. These issues together mean that future predictions of gene drive spread will likely involve high levels of uncertainty.

Environmental carrying capacity is another key factor that determines the spread and persistence of the simulated gene drive releases, as well as the wild-type population abundance. The introgression of the gene drive tended to initially follow seasonal patterns of carrying capacity driven by precipitation (Figure 5). Later, however, the importance of resistance gradually overcame the drive within 11 years despite the spatio-temporal variability of the population abundances and a relatively mild genetic load imposed by the construct. Resistance is a recognised challenge for gene drive systems (e.g., Hammond et al., 2017; Price et al., 2020; James et al., 2020) for which various counter-strategies have been proposed (Beaghton et al., 2017; Nash et al., 2019; Nolan, 2021; Wang et al., 2021). Our simulations assume resistance alleles arise with frequency *r* (Table 2) determined by the probability of cleavage (*k*_*c*_), the probability of nonhomologous repair (*k*_*j*_) and the probability that the nuclease gene becomes non-functional due to mutation of the target site during homologous repair (*k*_*n*_) (Beaghton et al., 2017). Our model, however, does not account for preexisting resistance in wild type populations caused by sequence variation in the target locus. Hence the (nonetheless rapid) progression of resistance in hybridising and spatially heterogeneous populations shown here could be underestimated, further emphasising the importance of managing drive resistance.

Our characterisation of environmental carrying capacities and abundance for the wild-type populations can accommodate alternative functional forms and parametrisation. Moreover, the functional forms and parametrisation of carrying capacity may be expected to change with time as climate (Kelly-Hope et al., 2009; Afrane et al., 2012) and land use (Keating et al., 2004; Munga et al., 2009) changes. The current lack of quantitative, species-specific, data on mortality and dispersal within the *An. gambiae* s.l. complex, however, limits our ability to parametrise relationships such as the larval carrying capacity (Equation 2.9) in a species-specific fashion (North and Godfray, 2018), beyond a few, spatially limited, empirical studies (e.g. Magombedze et al., 2018). We anticipate that as data from entomological surveys, coupled with species differentiation through genetic methods (e.g., *Anopheles gambiae* 1000 Genomes Consortium, 2017; Clarkson et al., 2020; Giraldo-Calderón et al., 2015) is increasingly centralised, then more detailed parametrisations will become possible at the subspecies level.

A lack of data also constrains our ability to compare alternative hypotheses of how *An. gambiae* s.l. populations persist in marginal habitat zones such as the Sahel, where observed patterns seasonal abundance can be explained by either aestivation, permanence of larval microhabitats (i.e., non-zero carrying capacities during the dry season) or long range dispersal (North and Godfray, 2018). Although persistence in the Sahel can be explained without an explicit aestivation model (see Figures 3, 5), aestivation may nevertheless be an important life history strategy for some species within the *An. gambiae* s.l. complex in the Sahel (Lehmann et al., 2017). As for wind-driven dispersal, the relative importance of aestivation suffers from limited data.

The process model allows for the scenario-based testing of genetic and conventional vector control strategies. For genetic vector control, the sex-differentiated compartment model allows for both population replacement and population suppression gene drives. For the latter, the relative frequency of males and females may be an important component of the gene drive system (Kyrou et al., 2018) and also for sex-biased, self-limiting forms of genetic vector control strategies (Windbichler et al., 2008; Klein et al., 2012; Galizi et al., 2014); the model has the flexibility to accommodate sex bias from either maternal or paternal descent (Section 2.1.3). Conventional control strategies that target adult females or larval stage mosquitoes can be accommodated by increasing mortality rates in locations and times dependent on the intervention scenario (sensu Magombedze et al., 2018), or by reducing carrying capacity for interventions where larval habitat is removed.

Vector control is a key component in strategies developed to combat mosquito-borne and vector borne diseases (Wilson et al., 2020). The deployment of vector control strategies, whether genetic or conventional, into hybridising spatially heterogeneous populations will require the development of spatially explicit models. These models should be constructed at a commensurate spatial and temporal scale of the intervention and include the possibility of resistance, which is not only a feature of genetic methods such as gene drive systems but is also an expected development for conventional strategies such as insecticide applications (Achee et al., 2019; Nolan, 2021). Numerical simulation based scenario assessments can be used to investigate alternative ecological hypotheses in concert with proposed vector control intervention strategies. The spatio-temporal model developed here shows the importance of resistance, vertical gene transfer among hybridising subspecies, long range dispersal, spatio-temporal variability in larval mosquito habitats and deployment strategies at the continental scale.

## Supporting information

Animation (9 hours, A. coluzzii)

Animation (2 hours, A. gambiae s.s.)

Animation (2 hours, A. coluzzii)

Animation (9 hours, A. gambiae s.s.)

## Acknowledgments

The authors gratefully acknowledge the support of CSIRO Data61, CSIRO Health and Biosecurity and the Foundation for the National Institutes of Health. We would also like to thank Roslyn Hickson and Mathieu Legros for their thoughtful comments.

## Data availability

Illustrative animations are available: see Supplementary Materials Appendix S3.

## Supplementary Materials

### S1: Algorithm for placing introduction sites

As mentioned in the main text, we first select the cells in which at least 10,000 of either subspecies are available year-round on the African mainland. We then randomly select a set *Y* consisting of 5,000 of those cells as site candidates for ease of computation, discarding the others from consideration. We adapt an algorithm that maximises the probability of animals being captured by a set of traps spread across a domain using gradient descent.

We define a goodness of fit *G*(*X*) of a candidate set of 10 release sites *X* by:

- defining a kernel function *f* (*d*) = exp(− *d/*1000) where *d* is distance in km (analogous to probability of detection), then
- for each candidate site *y* ∈ *Y*, calculate 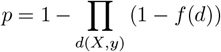 where *d*(*X, y*) is the set of distances between each point in *X* and the candidate *y* (analogous to the probability of detection by at least one site in *X*),
- taking *G*(*X*) as the sum of *p* over all the candidate sites (analogous to the expected number of candidate cells in *Y* detected by the sites in *X*).

For 20 iterations, we:

- initially select *X* to be 10 sites selected at random from *Y*, and
- calculate *G*(*X*) for this initial *X*.
- For each of the currently selected release sites *x* ∈ *X*:
  - pick the 20 spatially closest candidates to *x* and recalculate *G*(*X*) for each, temporarily replacing *x* with the nearby candidate in the set *X*; and
  - permanently replace *x* with the candidate that gives the highest *G*(*X*) in the set *X*, if higher than the current value.
- If the set *X* has changed after going through all of the release sites, then repeat the process.
- Otherwise, stop, outputting the final set *X* and final goodness of fit *G*(*X*).

Of the 20 output sets *X*, we then use the set that gave the highest value for *G*(*X*).

### S2: Neural network details

The inputs of the neural network were normalised by the mean and standard deviation of each predictor. The network was given three hidden layers of size 40, 20 and 10 (arrived at by experimentation; these sizes gave near-optimal validated errors: see Table 3); a rectified linear (ReLu) activation function for the hidden layers, and a sigmoid activation function for the output neuron to produce a probability of the predictors producing a mosquito of *An. Gambiae* s.s. as opposed to *An. coluzzii*. We use a cross-entropy loss function, such that the neural network effectively finds the maximum likelihood of the neuron weights given the observed subspecies at each site. The neural network is implemented using the Keras package (Chollet et al., 2017) in the R programming language (R Core Team, 2020). We use a 75:25 training-testing split on the data, using the DUPLEX algorithm (Snee, 1977) to ensure a spread of geographic locations for both training and testing sets. We then run the neural network for 200 epochs, and use the minimum validated loss to determine the number of epochs to subsequently run the model for.

For each tested set of predictors in the forward selection process, we train the neural network 500 times, running for 200 epochs each time. We calculate the validation loss after each epoch, take the minimum as our measure of goodness-of-fit for that trained network, and take the mean of the 500 validation losses obtained in this way. This is to account for variations in neural network effectiveness caused by the stochastic gradient descent process. Starting with an empty set of predictors, the predictor whose inclusion gives the best mean validation loss is accepted, until accepting new predictors no longer reliably decreases the validation loss.

We then train a neural network on the final selected set of predictors 100 times, taking the pointwise mean of the 100 runs as our final estimate of relative abundance at each location. This model averaging is performed in order to ensure that our final ensemble estimate is representative of the neural network modelling process as a whole: as the optimisation process is stochastic, a single network may differ substantially from another, especially where little data is available.

### S3: Illustrative animations

GIF animations of the simultaneous introduction of the construct at all sites are available in Supplementary Files: `wind9hF.ag.1.gif`, `wind9hF.ac.1.gif`, `wind2hF.ag.1.gif`, `wind2hF.ac.1.gif` show the animations for 9 hour wind advection for *Anopheles gambiae* s.s. and *Anopheles coluzzii*, and the 2 hour equivalents for both subspecies, respectively. The release points are marked as pink squares. Cells are colour coded: the amount of red in a cell represents the relative number of mosquitoes of the given subspecies with a wildtype *w* allele. Green and blue similarly represent construct *c* and resistance *r* respectively.

---

**Figure S1:**
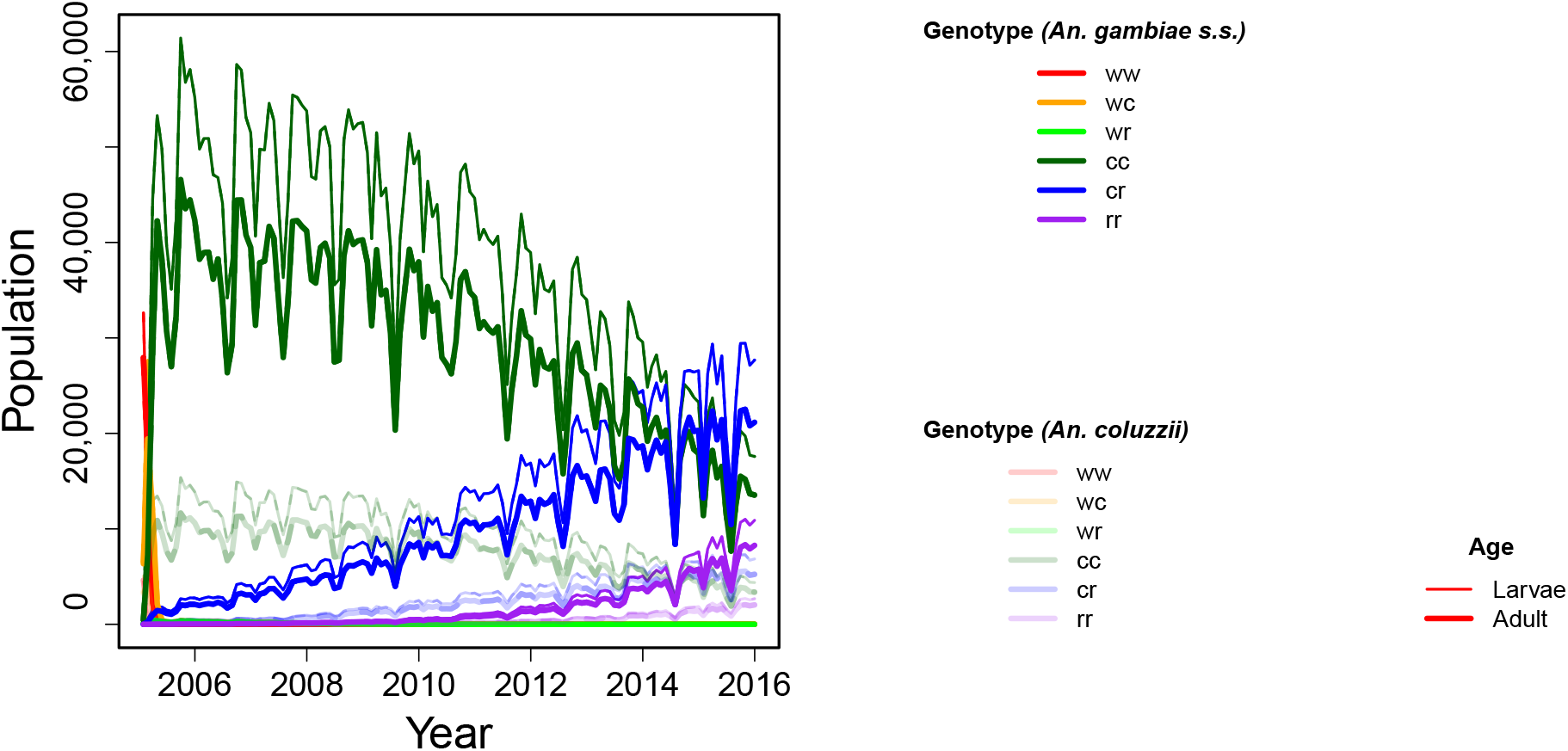
Time series plot of Site 6 **(as in Figure 5)** but with a closed population, i.e. no diffusion or advection. The colours correspond to genotype and the line thickness to age class.

**Figure S2:**
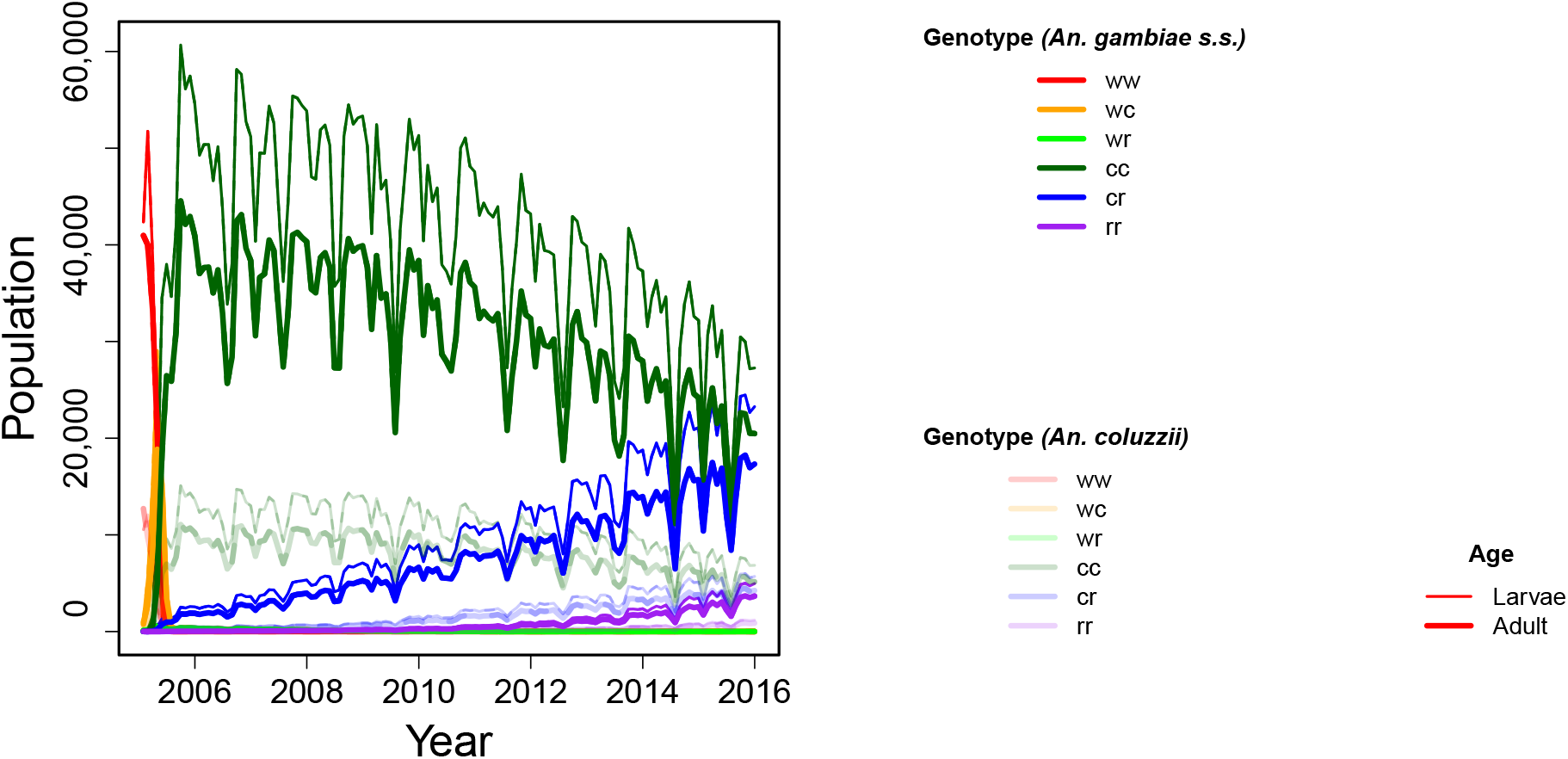
Time series plot of Site 6 **as in Figure S1** but with 6 age classes, keeping overall larval emergence period and mortality constant. This result is more closely analogous to a fixed larval emergence time than to the exponentially-distributed version in the main model. The first five age classes combined are plotted here under “larvae”. The colours correspond to genotype and the line thickness to age class.

**Figure S3:**
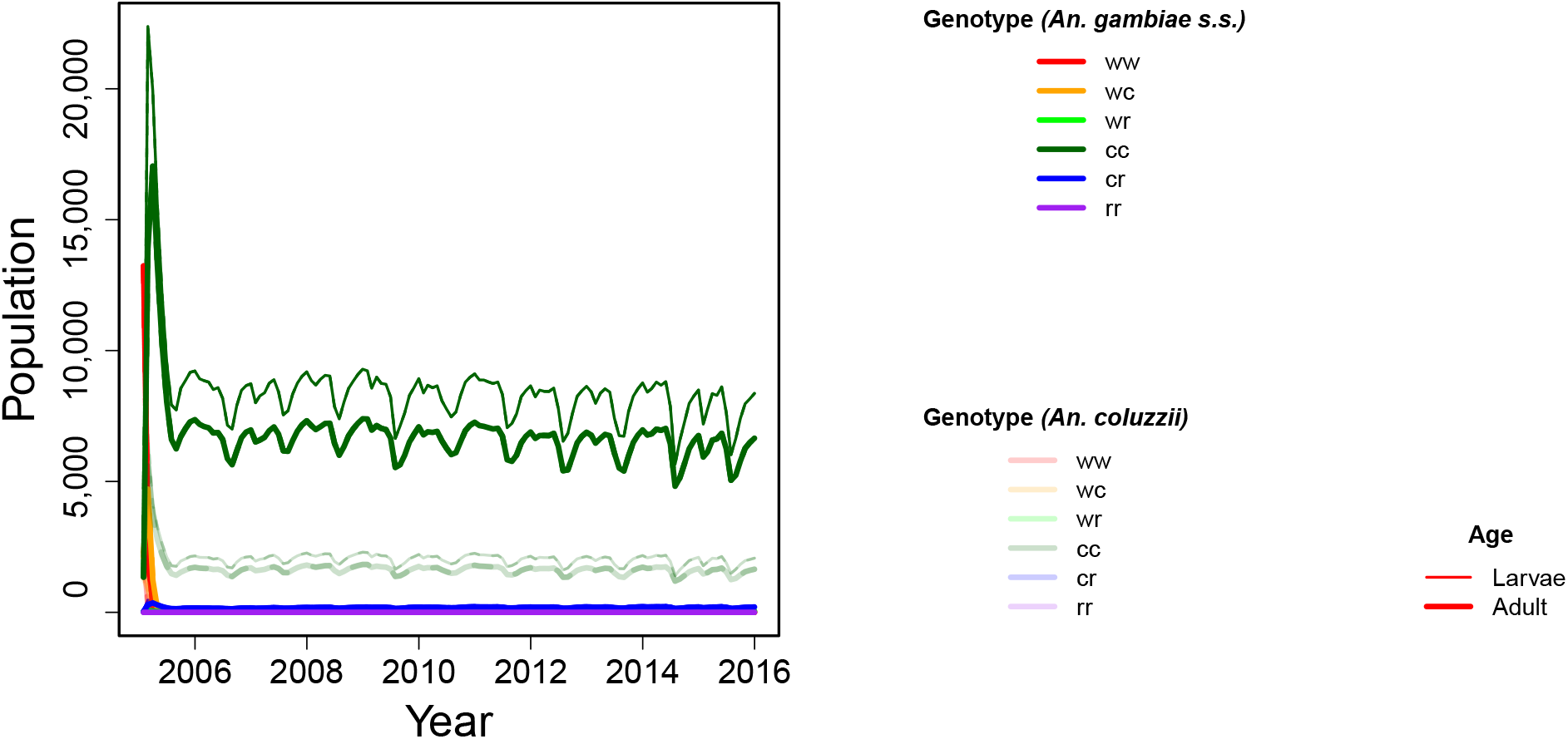
Time series plot of Site 6 **as in Figure S1** but with *R*(*g*_*M*_, *g*_*F*_) = 0.05 where *g*_*M*_ ≠ *ww* or *g*_*F*_ ≠ *ww*. The colours correspond to genotype and the line thickness to age class.

**Figure S4:**
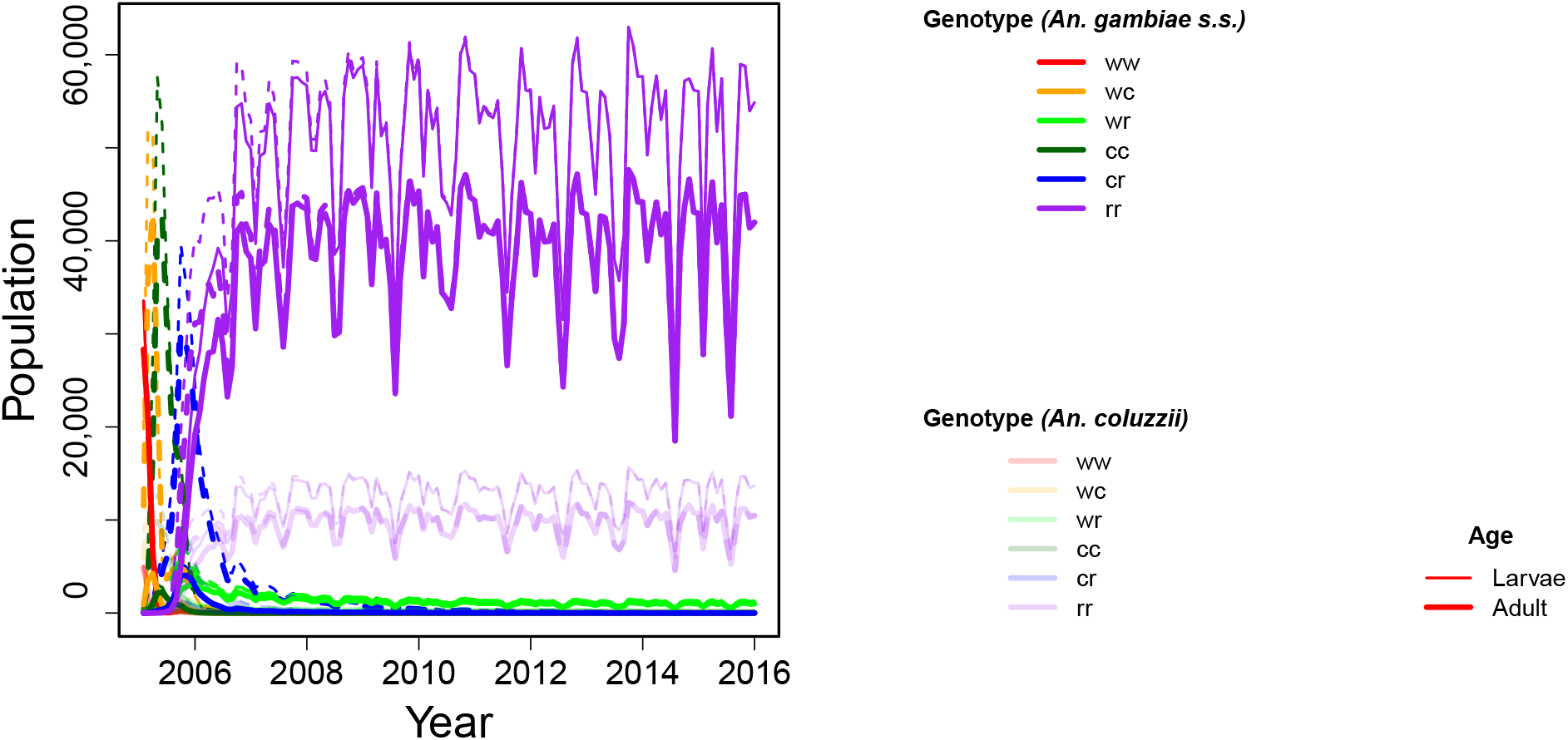
Time series plot of Site 6 **as in Figure S1** but with *p*(*g*_*M*_, *g*_*F*_, *s*) modified so that any presence of the genetic construct in the father (*wc, cc* or *cr*) results in a 95% male sex bias in offspring. Dashed lines represent males, solid lines females. The colours correspond to genotype and the line thickness to age class.

**Figure S5:**
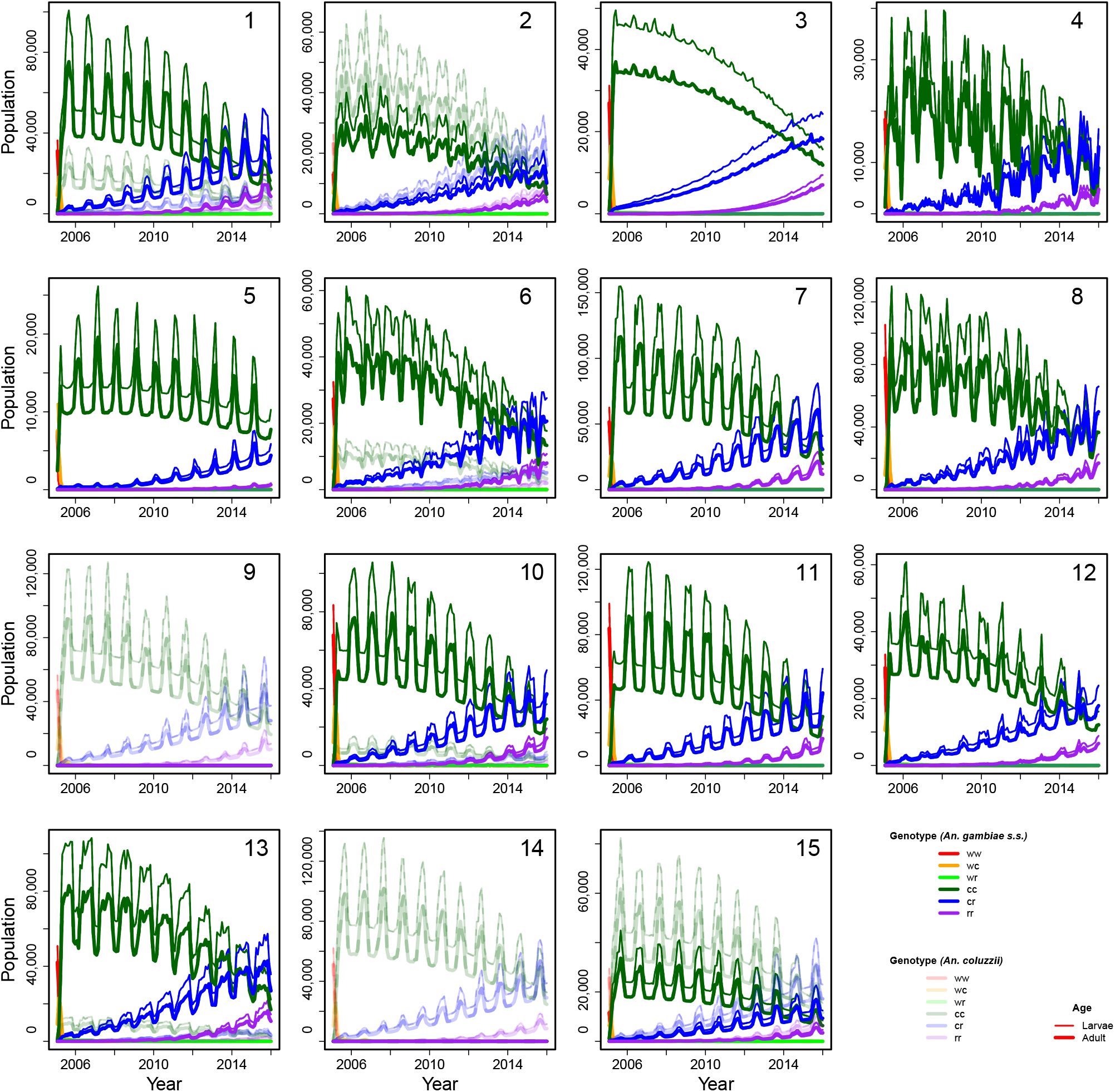
The time series abundance of male mosquitoes at each introduction point **as in Figure 5, but using 2 hours instead of 9 hours advection**, separated by species, genotype and age (female mosquitoes occur in identical numbers to males in this model). The colours correspond to genotype and the line thickness to age class.

**Figure S6:**
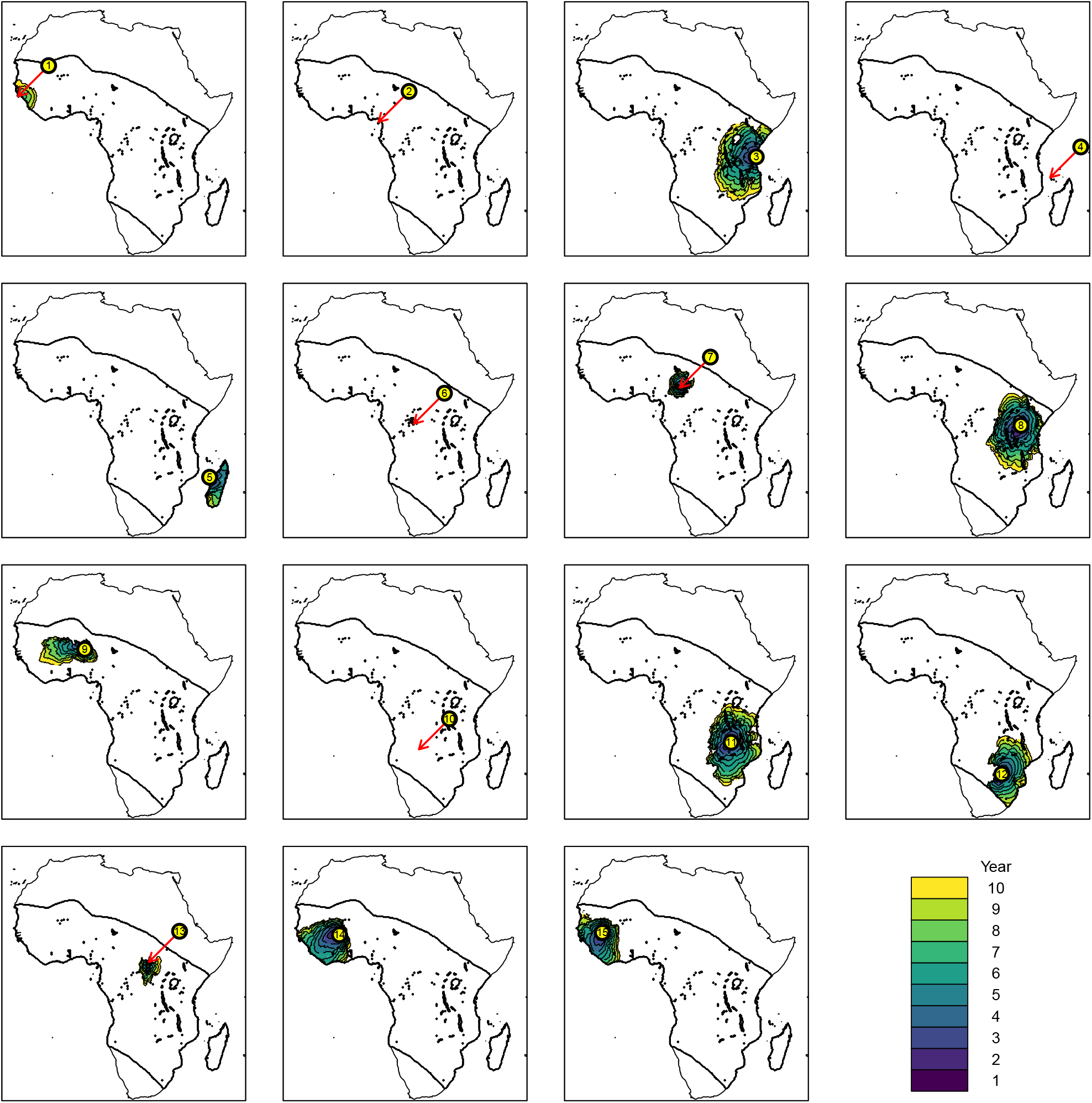
The invasion front of the construct at each introduction point **as in Figure 4, but using 2 hours instead of 9 hours advection**. A separate colour is given for each year. The island introductions are (1) the Bijagós islands (off Guinea-Bissau), (2) Bioko (off Cameroon), (3) Zanzibar (off Tanzania), (4) Comoros (off Mozambique) and (5) Madagascar.

**Figure S7:**
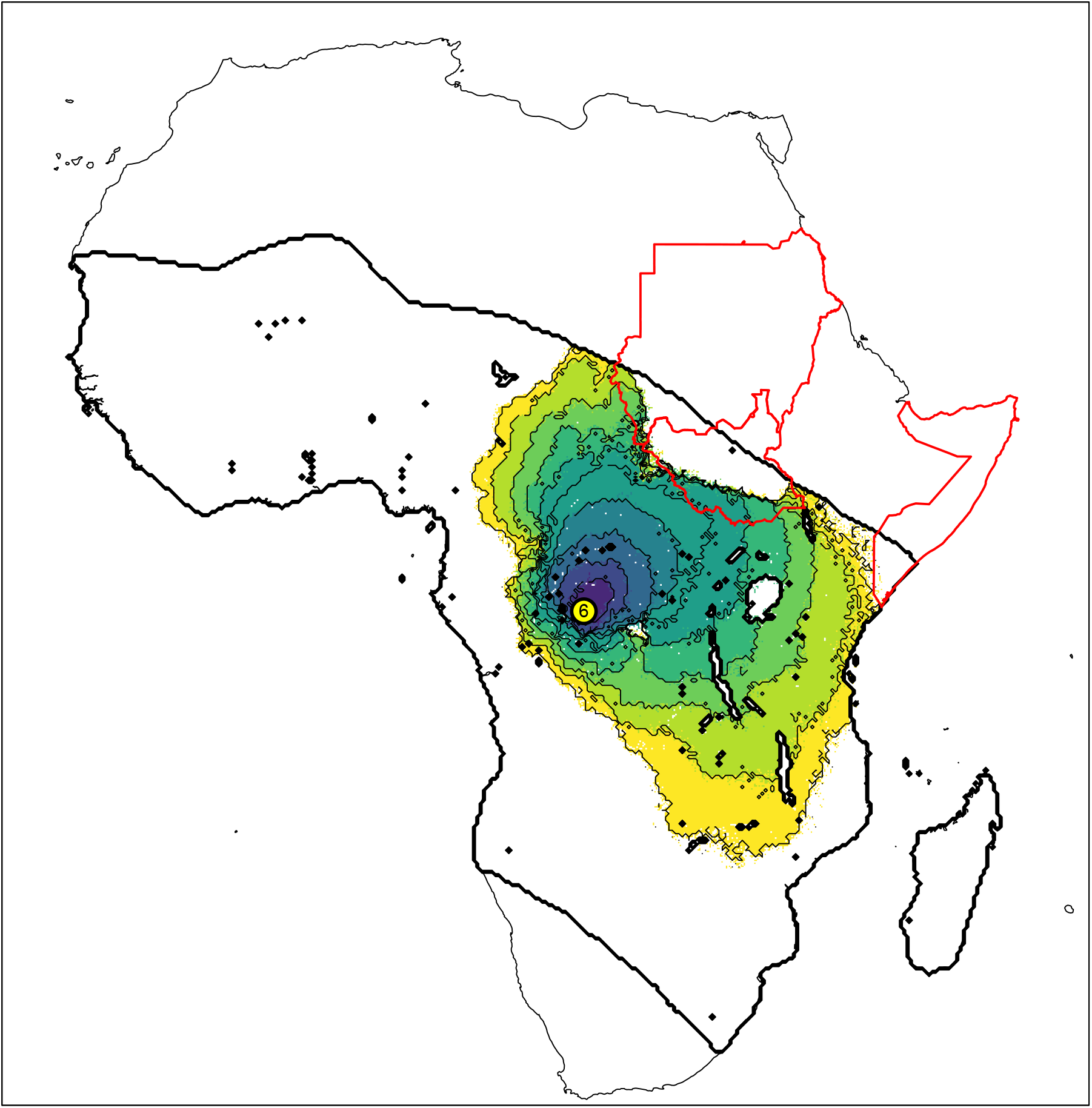
The invasion front of the construct as in Figure 4 for Site 6, but assuming human (and thus mosquito) absence in Sudan, South Sudan and Somalia (national boundaries given in red, as defined by the UN Office for the Coordination of Humanitarian Affairs).

