## Supplementary figures and images for "Spatial modelling for population replacement of mosquito vectors at continental scale"

### Animation (2 hours, A. coluzzii)

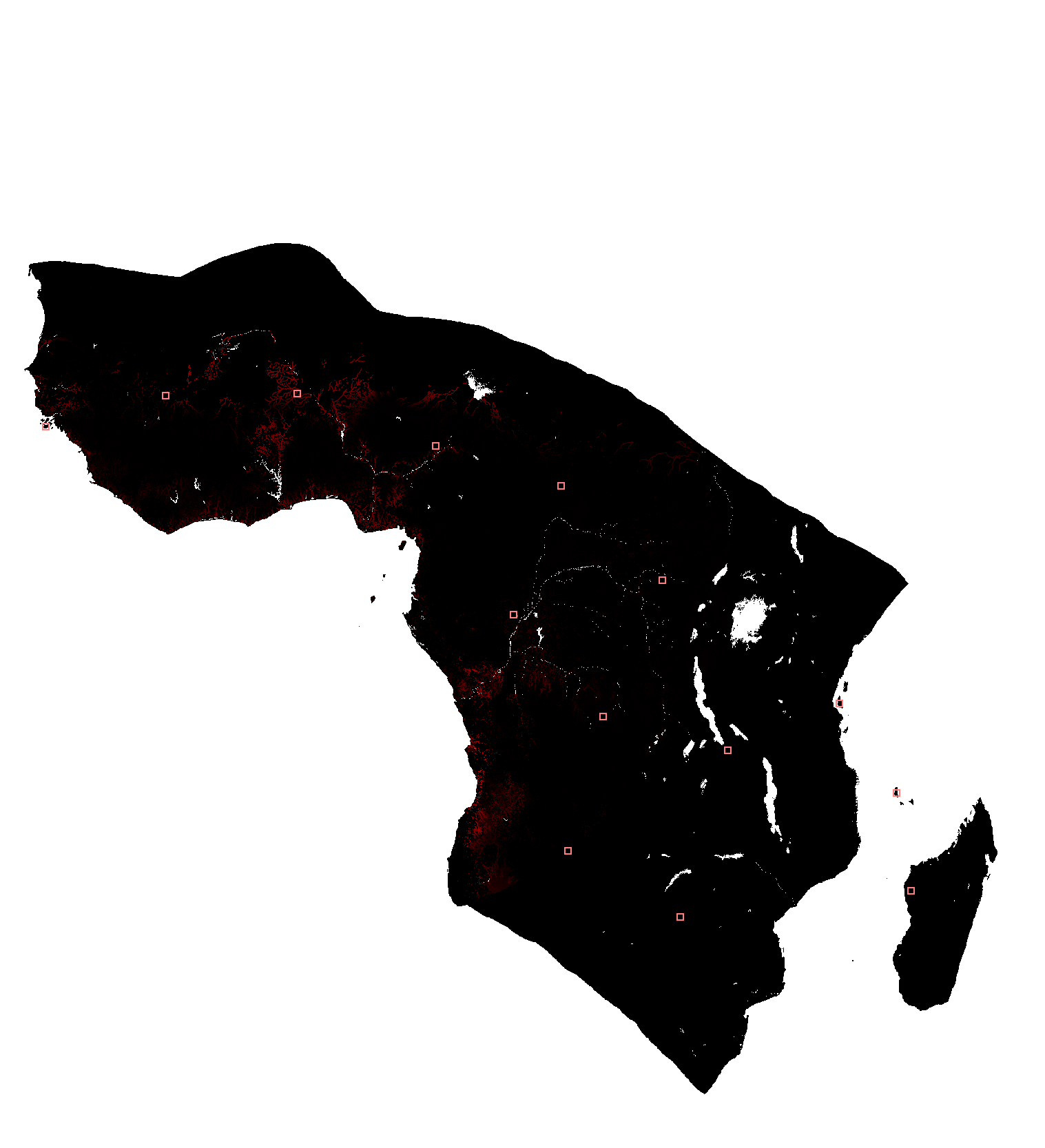

### Animation (2 hours, A. gambiae s.s.)

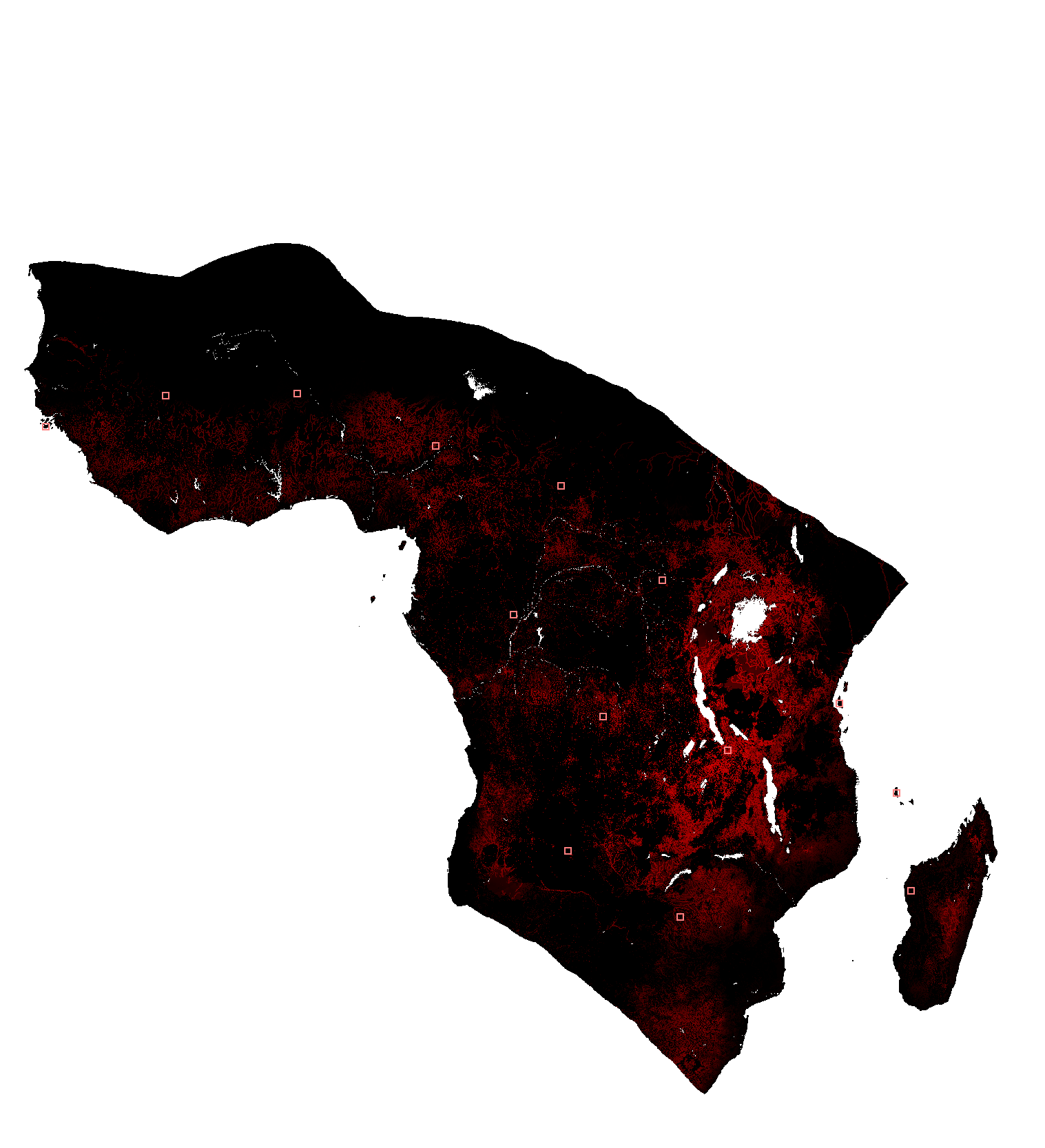

### Animation (9 hours, A. coluzzii)

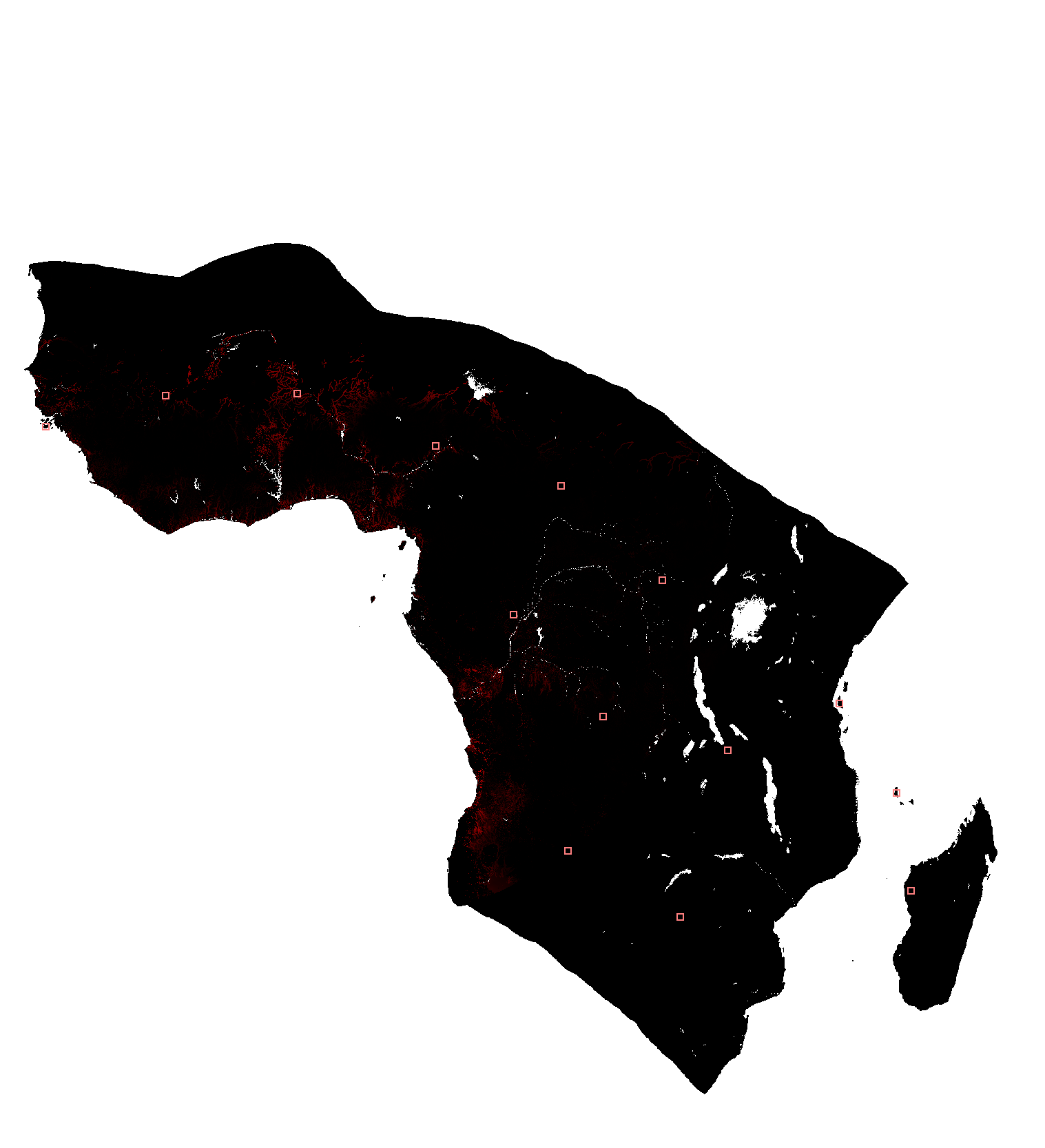

### Animation (9 hours, A. gambiae s.s.)

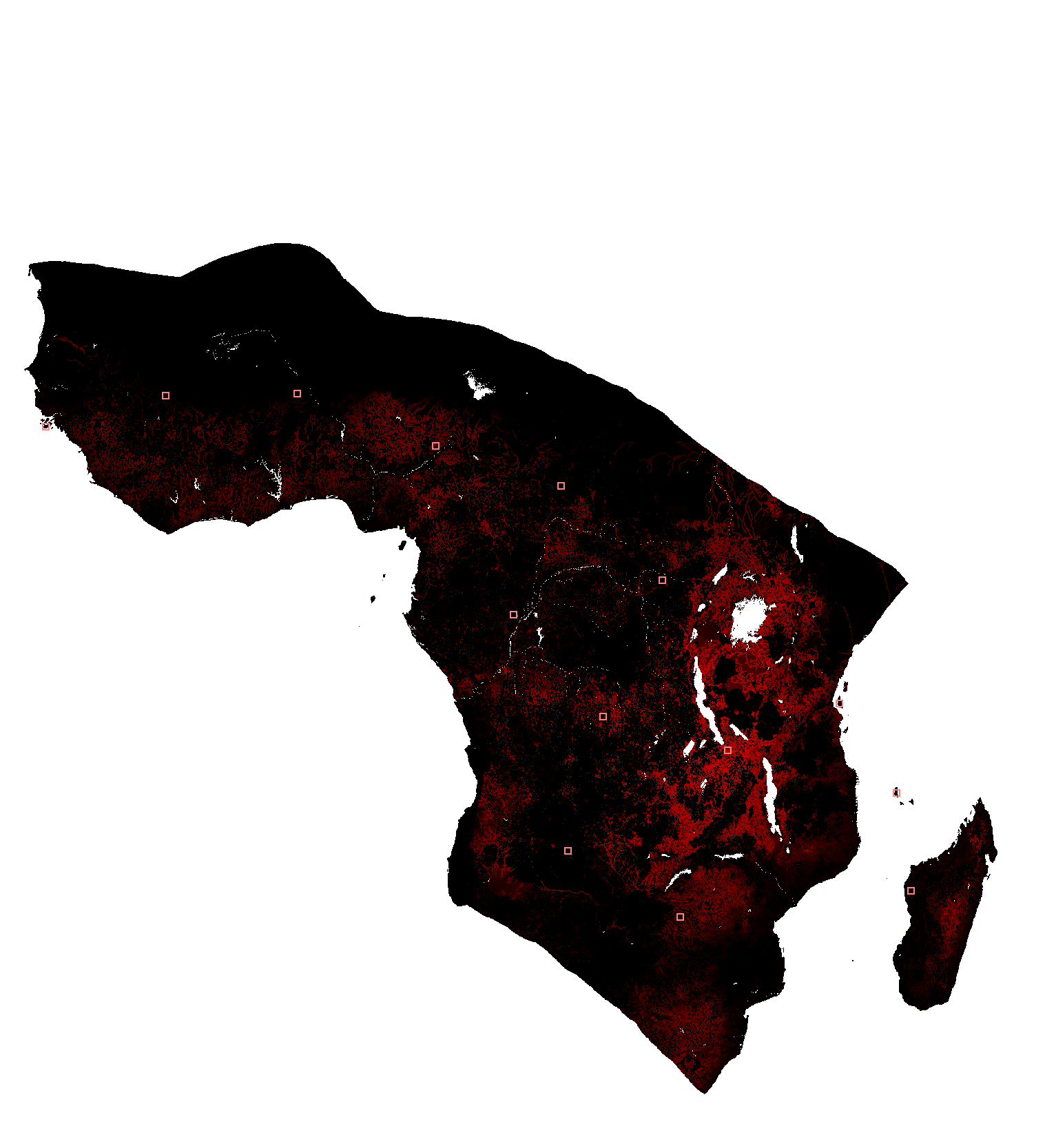
